# Structural effects and functional implications of phalloidin and jasplakinolide binding to actin filaments

**DOI:** 10.1101/794495

**Authors:** Sabrina Pospich, Felipe Merino, Stefan Raunser

## Abstract

Actin undergoes structural transitions during polymerization, ATP hydrolysis and subsequent release of inorganic phosphate. Several actin binding proteins sense specific states during this transition and can thus target different regions of the actin filament. Here we show in atomic detail that phalloidin, a mushroom toxin that is routinely used to stabilize and label actin filaments, suspends the structural changes in actin, likely influencing its interaction with actin binding proteins. Furthermore, high-resolution cryo-EM structures reveal structural rearrangements in F-actin upon inorganic phosphate release in phalloidin-stabilized filaments. We find that the effect of the sponge toxin jasplakinolide differs from the one of phalloidin, despite their overlapping binding site and similar interactions with the actin filament. Analysis of structural conformations of F-actin suggests that stabilizing agents trap states within the natural conformational space of actin.

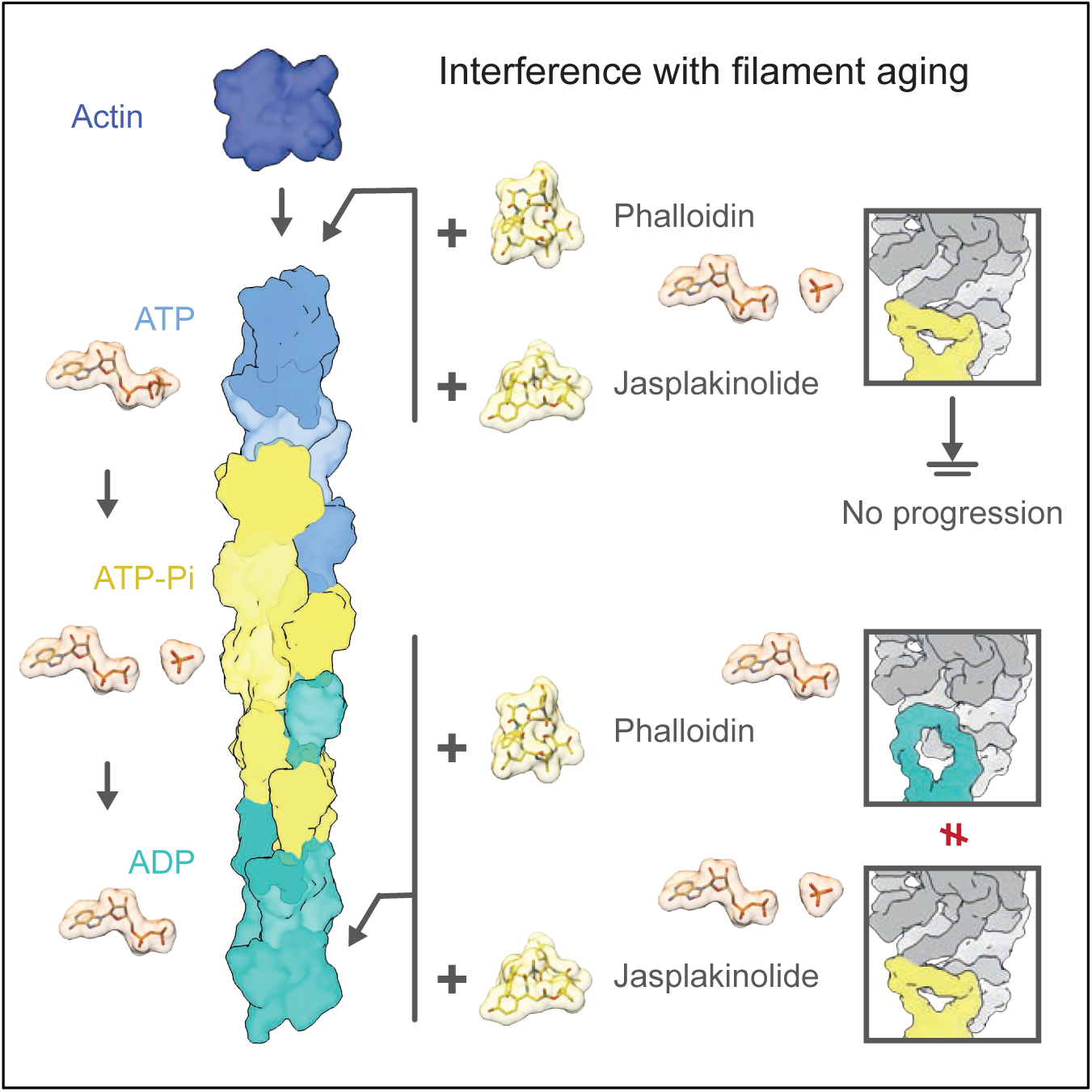

**Highlights:**

- Five high-resolution cryo-EM structures of stabilized filamentous actin
- Phalloidin traps different structural states depending on when it is added
- The effect of phalloidin and jasplakinolide on filamentous actin is not identical
- Both toxins likely interfere with the binding of proteins sensing F-actin’s nucleotide state

## Introduction

Actin is a highly conserved and abundant protein which is essential for many key processes including cell motility, muscle contraction and vesicle trafficking (Pollard and Cooper, 2009). The 42 kDa protein consists of four subdomains (SDs) enclosing the central metal-nucleotide binding site, which is occupied by Mg^2+^-ATP in globular actin (G-actin) (Kabsch et al., 1990). Under physiological conditions, G-actin spontaneously polymerizes into a double stranded right-handed helix (F-actin) that is stabilized by interactions between subunits of the same strand (intra-strand interactions) and across strands (inter-strand interactions) (Holmes et al., 1990).

During polymerization, G-actin undergoes a conformational change, resulting in the flatter appearance of actin subunits in F-actin (Oda et al., 2009) and a rotation of His161 into its catalytic position (Merino et al., 2018) enhancing the actin-ATPase activity by four orders of magnitude (Blanchoin and Pollard, 2002). ATP hydrolysis is followed by the slow release of inorganic phosphate (P_i_), which leaves ADP in the nucleotide-binding pocket of ‘aged’ F-actin. Thus, F-actin subunits pass through at least three different conformational states, namely the ATP state, the ADP-P_i_ state and the ADP state (Carlier and Pantaloni, 1986). The final ADP state is the least stable of the three states (Merino et al., 2018; Pollard, 1986). Due to the faster addition of ATP-bound G-actin to the barbed end, F-actin displays a characteristic nucleotide gradient along the filament, which marks the local age of the polymer.

We recently showed that nucleotide hydrolysis and P_i_ release in F-actin modulate the conformation of the periphery of the filament (Merino et al., 2018). In particular, we identified two distinct conformations which we named open and closed D-loop states based on substantial structural differences in the DNase-I-binding loop (D-loop). The open D-loop state is characterized by an outwards bent D-loop and an extended C-terminus. In the closed conformation, the D-loop is folded in and the C-terminus adopts a helical fold. We found that the two D-loop states coexist in the ATP-like and ADP-P_i_ states of F-actin whereas in F-actin-ADP, the D-loop is only found in its closed state. Another study also reported changes within the D-loop/C-terminus interface upon P_i_ release (Chou and Pollard, 2019). In general, this highlights the importance of this site, which is likely sensed by actin-binding proteins (ABP) like cofilin and coronin (Cai et al., 2007; Ge et al., 2014; Merino et al., 2018; Suarez et al., 2011).

Traditionally, natural compounds, such as phalloidin or jasplakinolide have been used to stabilize and study actin filaments. Phalloidin is a bicyclic heptapeptide originally isolated from the death cap mushroom *Amanita phalloides* (Lynen and U. Wieland, 1938) and well known for its specific, stochiometric binding to F-actin (Estes et al., 1981; T. Wieland and Govindan, 1974). Jasplakinolide (JASP), a cyclic depsipeptide originally isolated from the marine sponge *Jaspis johnstoni* (Crews et al., 1986), shares this characteristics and competitively inhibits the binding of phalloidin to F-actin (Bubb et al., 1994). Both toxins stabilize actin filaments and prevent depolymerization even under harsh conditions (Dancker et al., 1975; Lengsfeld et al., 1974). Under non-polymerizing conditions, they promote polymerization (Bubb et al., 1994; Estes et al., 1981). JASP has been reported to stabilize F-actin more effectively than phalloidin (Bubb et al., 2000; Visegrády et al., 2005; 2004). While both toxins do not affect ATP hydrolysis, they strongly delay the release of P_i_ (Barden et al., 1987; Dancker and Hess, 1990; Vig et al., 2011), suggesting a close correlation between filament stability and the presence of P_i_.

Over the years tremendous effort was put into the design and synthesis of phalloidin and JASP derivatives to enable specific labeling of actin filaments and to exploit their potential as therapeutics in anticancer or antimalaria therapy (Anderl et al., 2012; Holzinger, 2009; Lukinavičius et al., 2014; Terracciano et al., 2005; E. Wulf et al., 1979; Yao et al., 2019). Phalloidin is widely used to stabilize F-actin in electron microscopy studies and is a common marker for F-actin in fluorescence light microscopy when conjugated to fluorophores (Melak et al., 2017). Despite its advantage of cell permeability over phalloidin, JASP has only rarely been used for the same purpose.

First experiments using X-ray fiber diffraction, mutational studies, scanning transmission electron microscopy or computational docking (Belmont et al., 1999a; Drubin et al., 1993; Lorenz et al., 1993; Oda et al., 2005; Steinmetz et al., 1998), identified the binding site of phalloidin at the interface of three actin protomers, but failed to determine its exact position within the filament. Only recent high-resolution electron cryo microscopy (cryo-EM) studies revealed the exact interaction of phalloidin and JASP with F-actin (Iwamoto et al., 2018; Mentes et al., 2018; Merino et al., 2018; Pospich et al., 2017). Both toxins bind non-covalently to the same site consisting of three actin subunits from both strands and stabilize the filament by extensive hydrophobic interactions. Additional hydrogen bonds have been proposed to stabilize the interaction with phalloidin (Mentes et al., 2018).

In our latest study, we found that JASP traps the open D-loop conformation and inhibits P_i_ release when added immediately before polymerization of either G-actin-ATP or G-actin-ADP (Merino et al., 2018). Interestingly, the open D-loop state has not been observed in the structures of phalloidin-stabilized F-actin in complex with myosin or filamin A, which could be due to differences between JASP and phalloidin or the binding of myosin and filamin A (Iwamoto et al., 2018; Mentes et al., 2018). Considering this, the question arises if JASP and phalloidin act indeed in the same way on F-actin as presumed hitherto based on their biochemical properties.

In this work we systematically studied the effect of phalloidin on the structure of F-actin and compared it to the effect JASP has on actin filaments using cryo-EM. We found that both stabilizing agents interfere with the natural aging process of F-actin, thus camouflaging the nucleotide state of F-actin that in turn cannot be sensed by ABPs anymore. Furthermore, we describe structural changes upon P_i_ release and their coupling in phalloidin-stabilized F-actin. Principle component analysis suggests that the structural conformations trapped in toxin-stabilized filaments are representative for conformational states of F-actin in general.

## Results and Discussion

### The structural effect of phalloidin on F-actin

First, we determined the cryo-EM structure of filamentous rabbit skeletal muscle *α*-actin adding phalloidin immediately before polymerization (F-actin-PHD) to make sure that ATP complexed by actin was not hydrolyzed at the time phalloidin bound to F-actin. The average resolution of the reconstruction of ∼ 3.3 Å allowed the clear positioning of most side chains (Figure S1, Table 1). The structure of F-actin-PHD closely resembles the structure of F-actin-ADP-P_i_ JASP that was analogously prepared (Merino et al., 2018). The D-loop is well resolved and solely populates the open D-loop state (Figure 1A, Movie S1A-B), and the nucleotide binding pocket is occupied by ADP and P_i_ (Figure 2A, Movie S2A-B, Figure S2A). This is in line with previous reports where it was shown that JASP and phalloidin dramatically reduce the release of P_i_ from F-actin (Dancker and Hess, 1990; Vig et al., 2011). However, the density corresponding to P_i_ is weaker in comparison to F-actin-ADP-P_i_-JASP (Merino et al., 2018), indicating that the P_i_ affinity of F-actin-PHD is lower than in the JASP-bound F-actin. Since previous studies showed that JASP and phalloidin treatment results in similarly reduced P_i_ release rates (Vig et al., 2011), this shows that the k_on_ for P_i_ binding is higher in the case of F-actin-ADP-P_i_-JASP. Furthermore, phalloidin seems to interfere with the conformational coupling of the active site to the periphery considering that the low P_i_ occupancy in F-actin-PHD does not result in a mixture of conformational states but traps F-actin solely in the open D-loop state.

**Table 1.** Data collection, refinement, and model building statistics of phalloidin data sets.

|  | F-actin-PHD | F-actin-PHD-aged | F-actin-PHD-Alexa |
| --- | --- | --- | --- |
| **Microscopy** |  |  |  |
| Microscope | Titan Krios (X-FEG, Cs-corrected) |  |  |
| Voltage [kV] | 300 |  |  |
| Defocus range [$\mu\text{m}$] | 0.6 – 2.8 | 0.7 – 2.8 | 0.4 – 2.6 |
| Camera | Falcon II linear | Falcon III linear | Falcon III linear |
| Pixel size [$\text{\AA}$] | 1.14 | 1.12 | 1.12 |
| Total electron dose [$\text{e}/\text{\AA}^2$] | 71 | 87 | 92 |
| Exposure time [s] | 2.1 | 1.5 | 1.5 |
| Frames per movie | 40 | 35 | 35 |
| Number of images<sup>a</sup> | 2,041<br>(3,425) | 2,205<br>(5,213) | 3,747<br>(5,221) |
| **3-D refinement statistics** |  |  |  |
| Number of helical segments<sup>a</sup> | 405,264<br>(598,021) | 513,783<br>(1,209,605) | 719,053<br>(905,946) |
| Electron dose final refinement [$\text{e}/\text{\AA}^2$] | 14 | 15 | 16 |
| Resolution [$\text{\AA}$]<sup>b</sup> | 3.3 | 3.7 | 3.6 |
| Map sharpening factor [$\text{\AA}^2$] | -86 | -128 | -126 |
| **Measured helical symmetry<sup>c</sup>** |  |  |  |
| Rise [$\text{\AA}$] | 27.59±0.02 | 27.56±0.03 | 27.53±0.02 |
| Helical twist [$^\circ$] | 166.9±0.1 | 166.9±0.1 | 166.9±0.1 |
| **Atomic model statistics** |  |  |  |
| Non-hydrogen atoms | 14,915 | 14,915 | 14,590 |
| Rosetta FSC-Work/FSC-Free | 0.574/0.565 | 0.586/0.584 | 0.583/0.582 |
| Molprobity score | 1.37 | 1.54 | 1.26 |
| Clashscore | 2.32 | 4.70 | 3.66 |
| EMRinger score | 3.37 | 2.85 | 2.96 |
| Bond RMSD [$\text{\AA}$] | 0.017 | 0.018 | 0.018 |
| Angle RMSD [$^\circ$] | 1.42 | 1.48 | 1.43 |
| Poor rotamers [%] | 0 | 0 | 0 |
| Favored rotamers [%] | 100 | 100 | 100 |
| Ramachandran favored [%] | 94.85 | 95.66 | 97.49 |
| Ramachandran outliers [%] | 0.00 | 0.00 | 0.00 |

<sup>a</sup>In parenthesis is the initial number of images or segments.<sup>b</sup>The resolution corresponds to a section of $\sim 120 \text{ \AA}$ at the center of the filaments<sup>c</sup>The helical parameters were estimated from the atomic model of five consecutive subunits independently fitted to the map as described before (Pospich et al., 2017). The small variations on the helical rise are within the margin of error of the pixel size determination. Therefore, the differences between reconstructions from different detectors could be due to technical differences.

**Figure 1.**
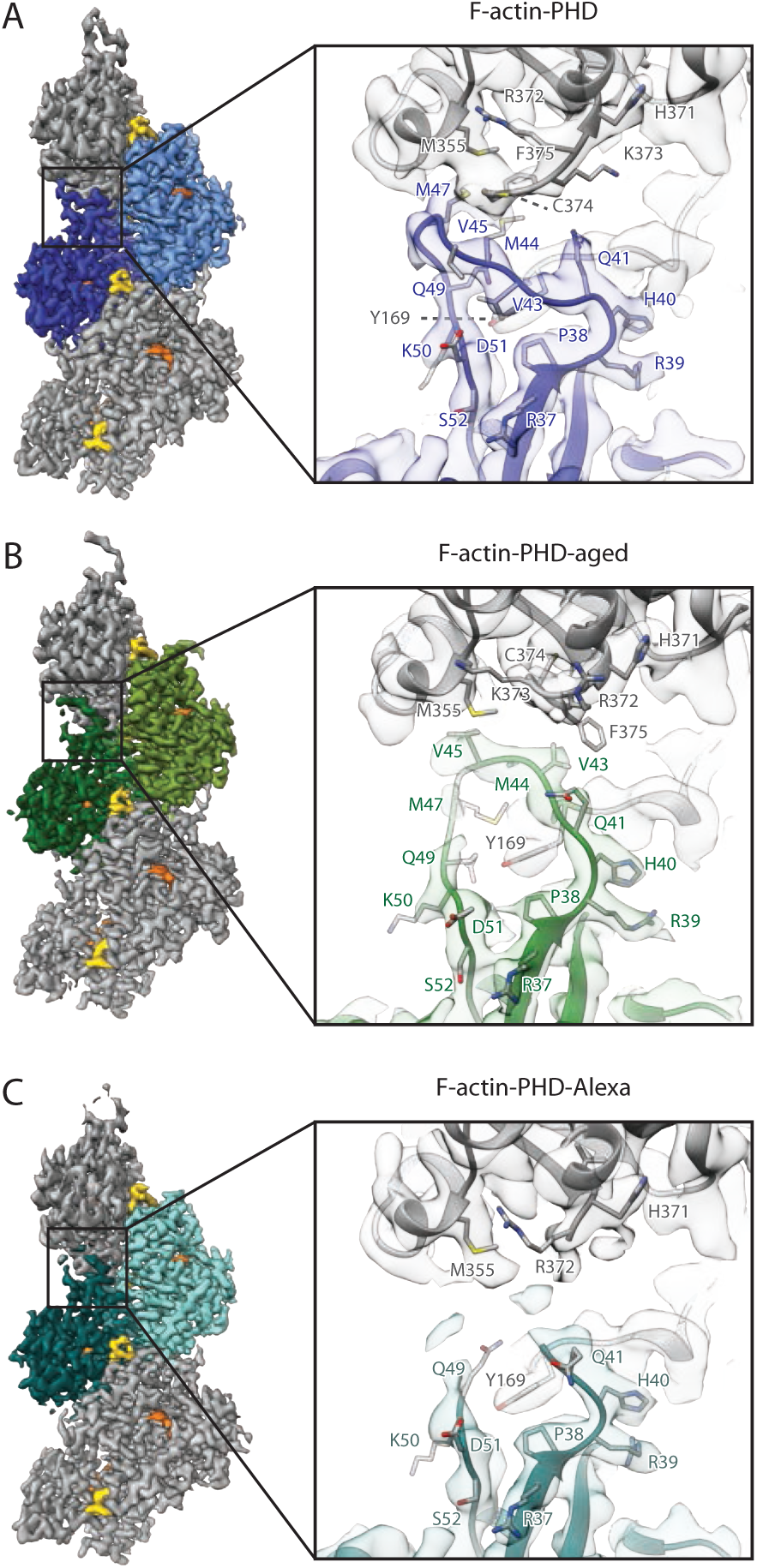
The intra-strand interfaces of phalloidin-stabilized F-actin. **(A-C)** Depiction of the intra-strand interfaces of **(A)** F-actin-PHD (shades of blue), **(B)** F-actin-PHD-aged (shades of green) and **(C)** F-actin-PHD-Alexa (shades of turquoise). While F-actin clearly adopts the open D-loop conformation in case of F-actin-PHD, the D-loop is fragmented and mixed for F-actin-PHD-Alexa and adopts a closed conformation accompanied by a helical-fold of the C-terminus in case of F-actin-PHD-aged. The tip of the D-loop and terminal part of the C-terminus of F-actin-PHD-Alexa were not included into the model due to the fragmentation of the density. Also see Movie S1A-C.

**Figure 2.**
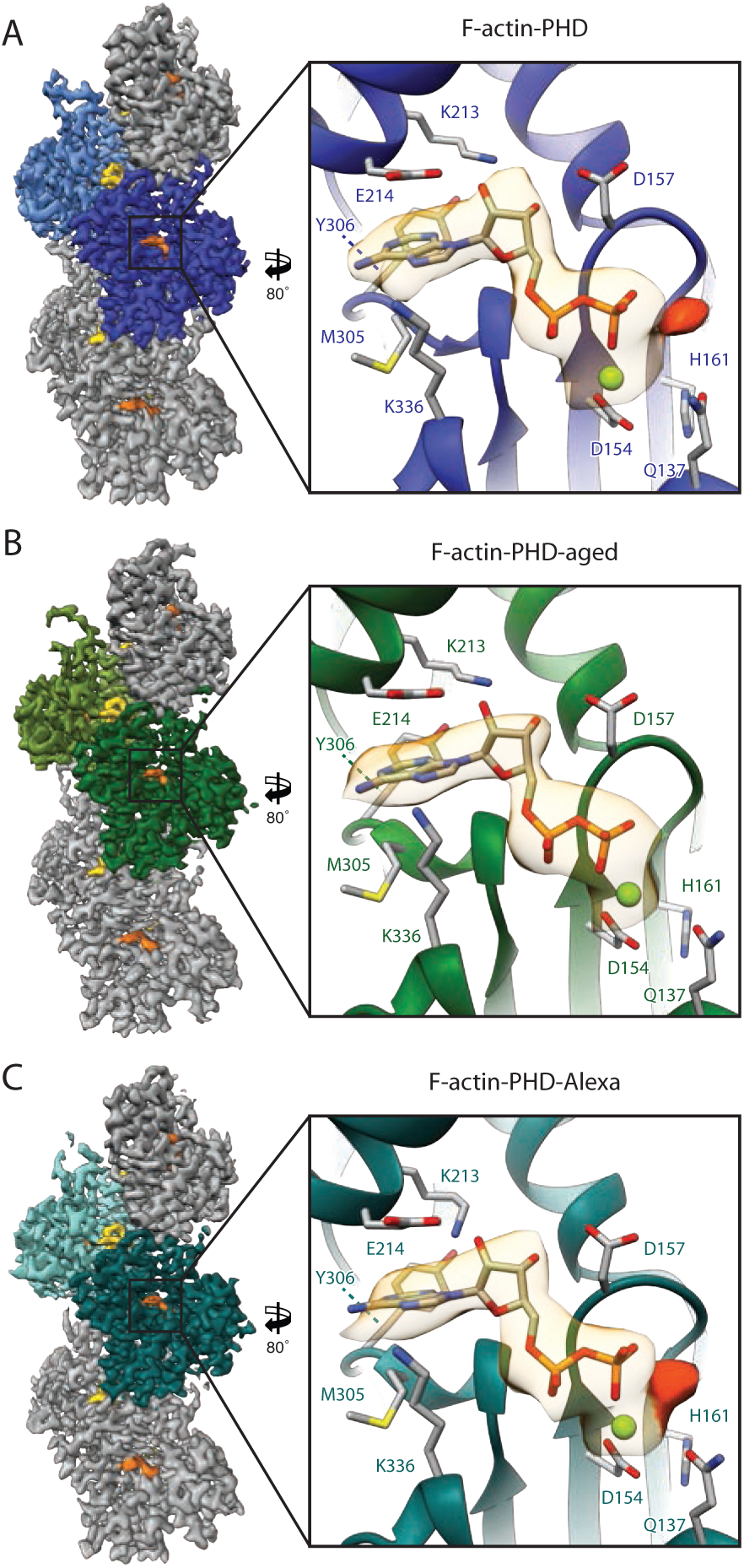
Nucleotides bound to phalloidin-stabilized F-actin. **(A-C)** Depiction of the central nucleotide binding cleft. Both F-actin-PHD and F-actin-PHD-Alexa have weak density for an inorganic phosphate (P_i_, highlighted by a darker shade of orange) which was not fully released after ATP hydrolysis. Due to its weak density, P_i_ was not included into the atomic models. After aging of filaments, one cannot find density corresponding to P_i_ anymore. Nucleotides and phalloidin are colored in orange and yellow, respectively. Also see Figure S2A-C and Movie S2A-C, for color-code see Figure 1.

To distinguish whether the closed D-loop conformation of F-actin-ADP (Ecken et al., 2015; Merino et al., 2018) is changed when adding phalloidin after polymerization and aging of the filament, we determined the ∼ 3.7 Å cryo-EM structure of F-actin in complex with phalloidin, in which the toxin was added one night after polymerization (F-actin-PHD-aged) (Figure S1, Table 1). This approach should guarantee that all ATP has been hydrolyzed and P_i_ has been completely released from the filament. The structure revealed that although phalloidin binds stoichiometrically to F-actin, the D-loop remains in the closed state (Figure 1B, Movie S1C-D). Expectedly, the active site only accommodates ADP without any trace of P_i_ (Figure 2B, Figure S2B, Movie S2C-D). This is in line with previous studies reporting that myosin-5 and myosin-6 can distinguish the ADP-P_i_ from the ADP state in phalloidin stabilized F-actin (Zimmermann et al., 2015). Considering our previous data indicating that the ABP coronin can sense the nucleotide state of F-actin via the D-loop conformation (Merino et al., 2018), we propose that myosin-5 and myosin-6 can read the age of F-actin in a similar manner by binding to the D-loop C-terminus interface (Gurel et al., 2017; S. F. Wulf et al., 2016).

At the moment, there is only one high-resolution cryo-EM structure of an actomyosin complex available that has been stabilized by phalloidin, namely the F-actin-myosin-1B complex (Mentes et al., 2018). In this structure, the D-loop is in the closed conformation and there is no density corresponding to P_i_, suggesting that phalloidin was added to aged filaments after polymerization. However, it is not described when phalloidin was added in the original paper. Therefore, no conclusions can be drawn regarding the specific recognition of the D-loop state by myosin. In general, as already stated by others (Zimmermann et al., 2015), this illustrates the necessity for future publications to describe in detail how phalloidin is used.

The binding site and structure of phalloidin in both F-actin-PHD and F-actin-PHD-aged closely resemble the ones described for the F-actin-myosin-1B structure (Mentes et al., 2018) (Figure 3A-F, Movie S3A-D). Phalloidin binds to the interface of three actin subunits from both strands, thereby strengthening intra- and inter-strand contacts. The interaction is primarily mediated through hydrophobic interactions (Figure S3A-C) complemented by additional putative hydrogen bonds. It has been previously suggested that Asp179, Gly197, Ser199, and Glu205 contribute in hydrogen bonding to phalloidin (Mentes et al., 2018). In addition, our models indicate the presence of putative hydrogen bonds also between phalloidin and Glu72 and Gln246. Interestingly, we found an extra density that likely corresponds to an ion bound to phalloidin in both maps (Figure 3B,E, Movie S3B,D). Although we cannot exclude calcium remaining from previous purification steps, the buffer composition suggests potassium and magnesium to be the primary candidates.

**Figure 3.**
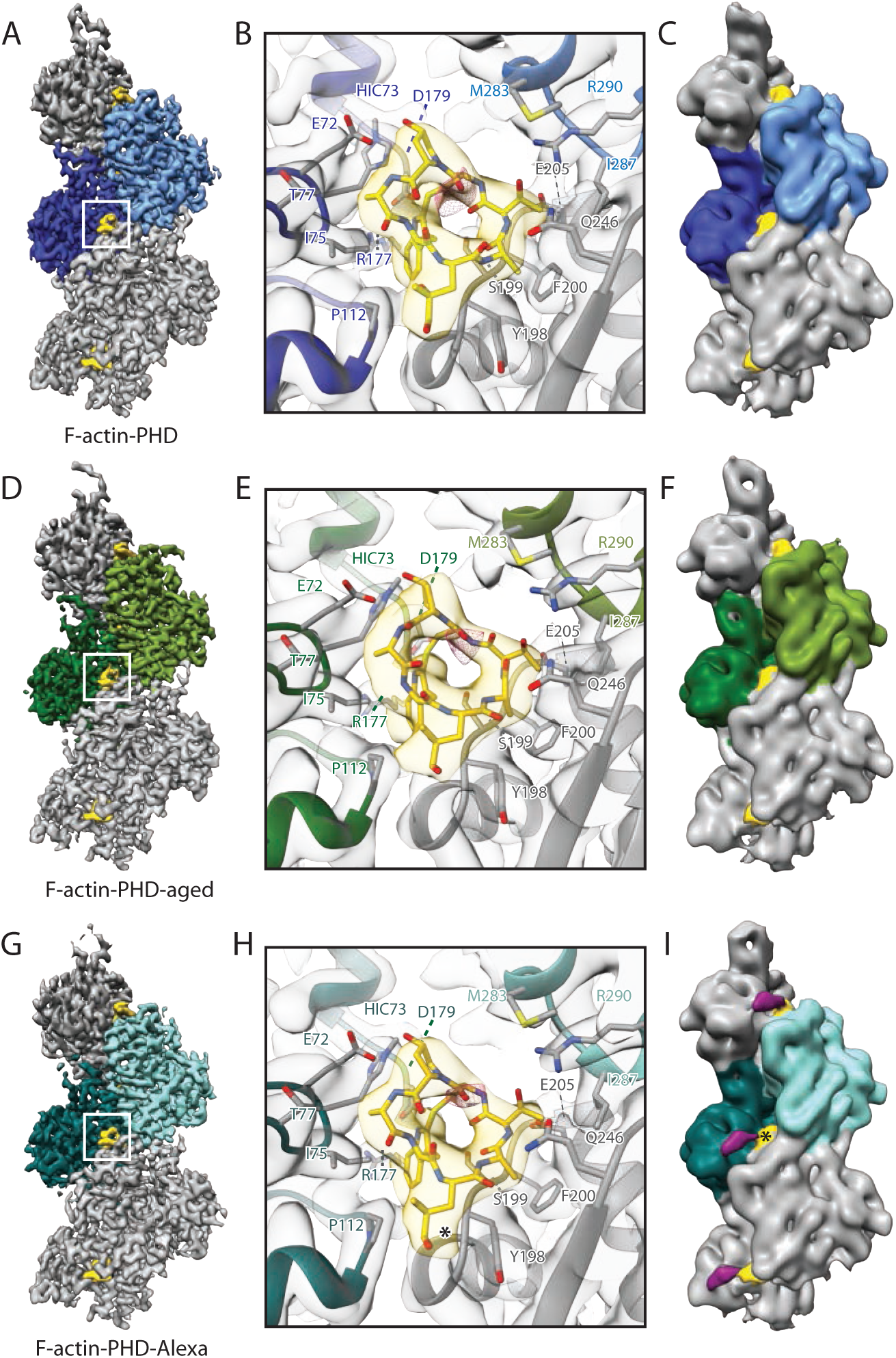
The phalloidin binding site. (**A-J**) Central binding sites illustrating the interaction of phalloidin (yellow) with three subunits from both strands. Extra density, possibly corresponding to an ion (pink), is present in all three structures. The Alexa Flour 546 dye (violet) within the F-actin-PHD-Alexa structure is not visible in the sharpened map filtered to local resolution **(G)**, but becomes visible in the unsharpened map filtered to 8 Å resolution **(I)**, suggesting pronounced flexibility. The position where Alexa Flour 546 is linked to phalloidin is marked with asterisks in **(H, I)**. Also see Movie S3A-C and Figure S3, for color-code see Figure 1.

### Differences in the effect of JASP compared to phalloidin

While the D-loop C-terminus interface adopts the closed conformation in the F-actin-PHD-aged structure, it is in the open state in our recent structure of F-actin-ADP copolymerized with JASP (Merino et al., 2018). To test whether this is due to the timepoint of addition of the respective toxin, we reconstituted F-actin-JASP in the same way as we have done with F-actin-PHD-aged, i.e. we added JASP after overnight ageing of F-actin (F-actin-JASP-aged). Using this approach and considering the similar properties of the toxins, we expected the filaments to be in the closed D-loop state with solely ADP in the nucleotide-binding pocket. However, the cryo-EM structure of F-actin-JASP-aged, which reached an average resolution of ∼ 3.7 Å (Figure S1, Table 2), differs significantly from our expectations.

**Table 2.** Data collection, refinement, and model building statistics of JASP data sets.

|  | F-actin-JASP-aged 1 | F-actin-JASP-aged 2 |
| --- | --- | --- |
| **Microscopy** |  |  |
| Microscope | Titan Krios (X-FEG, Cs-corrected) | Titan Krios (X-FEG, Cs 2.7 mm) |
| Voltage [kV] | 300 | 300 |
| Defocus range [$\mu\text{m}$] | 1.0 – 3.0 | 0.5 – 3.0 |
| Camera | Falcon III linear | K2 super-resolution |
| Pixel size [$\text{\AA}$] | 1.12 | 0.525 / 1.05<sup>a</sup> |
| Total electron dose [$\text{e}/\text{\AA}^2$] | 88 | 78 |
| Exposure time [s] | 2.1 | 15 |
| Frames per movie | 35 | 40 |
| Number of images<sup>b</sup> | 4,314<br>(5,335) | 1,994<br>(2,856) |
| **3-D refinement statistics** |  |  |
| Number of helical segments<sup>b</sup> | 352,285<br>(1,237,583) | 336,783<br>(467,060) |
| Electron dose final refinement [$\text{e}/\text{\AA}^2$] | 15 | Polished particles |
| Resolution [$\text{\AA}$]<sup>c</sup> | 3.7 | 3.1 |
| Map sharpening factor [$\text{\AA}^2$] | -116 | -48 |
| **Measured helical symmetry<sup>d</sup>** |  |  |
| Rise [$\text{\AA}$] | 27.61±0.02 | 27.67±0.02 |
| Helical twist [$^\circ$] | 166.8±0.1 | 166.7±0.1 |
| **Atomic model statistics** |  |  |
| Non-hydrogen atoms | 14,910 | 14,910 |
| Rosetta FSC-Work/FSC-Free | 0.598/0.598 | 0.622/0.623 |
| Molprobity score | 1.06 | 0.97 |
| Clashscore | 2.69 | 2.05 |
| EMRinger score | 2.80 | 3.58 |
| Bond RMSD [$\text{\AA}$] | 0.037 | 0.035 |
| Angle RMSD [$^\circ$] | 1.70 | 1.71 |
| Poor rotamers [%] | 0 | 0 |
| Favored rotamers [%] | 100 | 100 |
| Ramachandran favored [%] | 99.46 | 99.19 |
| Ramachandran outliers [%] | 0.00 | 0.00 |

<sup>a</sup> Pixel size after 2x binning used for processing<sup>b</sup> In parenthesis is the initial number of images or segments.<sup>c</sup> The resolution corresponds to a section of $\sim 120 \text{ \AA}$ at the center of the filaments<sup>d</sup> The helical parameters were estimated from the atomic model of five consecutive subunits independently fitted to the map as described before (Pospich et al., 2017). The small variations on the helical rise are within the margin of error of the pixel size determination. Therefore, the differences between reconstructions from different detectors could be due to technical differences.

First, the D-loop of F-actin-JASP-aged populates the open state to a large extent (weak density for the closed state, Figure 4A, Movie S1G-H), resembling our previous F-actin-ADP-P_i_-JASP structure (Merino et al., 2018). Second, the map includes strong density for P_i_ with an occupancy similar to ADP (Figure 5A, Movie S2G-H). This is surprising, since the P_i_ should be completely released similar to F-actin-PHD-aged (Figure 2B, Movie S2C-D). However, as for F-actin-PHD-aged, we did not remove the free P_i_ from the buffer but only reduced its concentration by pelleting the filaments and careful washing of the pellet for several times with phosphate-free buffer (see Methods). Therefore, the prominent difference regarding the presence of P_i_ in the nucleotide binding site suggests that JASP considerably increases the binding affinity of F-actin for P_i_ in comparison to phalloidin. This likely results in the remaining free P_i_ to be reincorporated into the active site.

**Figure 4.**
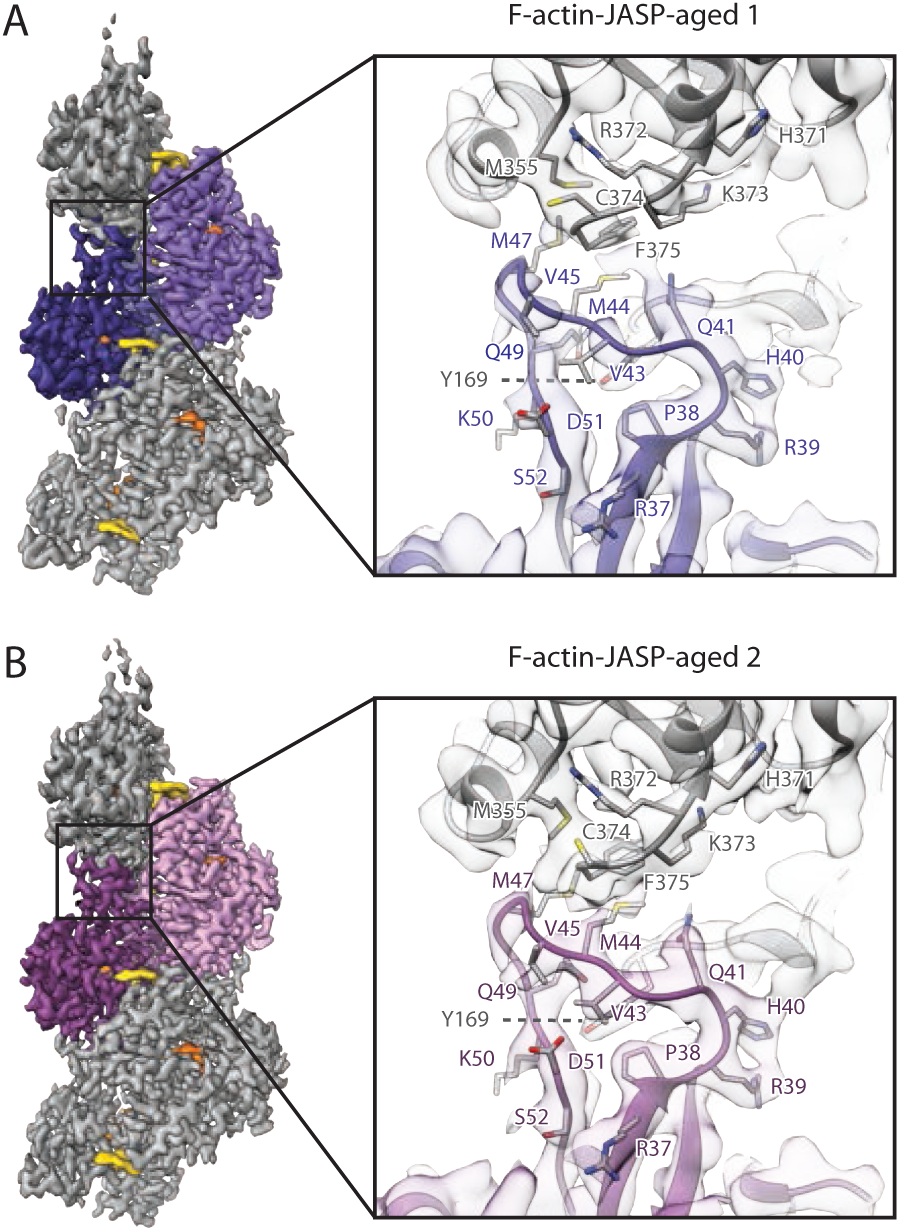
The intra-strand interfaces of JASP-stabilized F-actin. **(A-B)** Depiction of the central intra-strand interface made up by the D-loop and C-terminus of **(A)** F-actin-JASP-aged 1 (shades of purple) and **(B)** F-actin-JASP-aged 2 (shades of lavender). While F-actin-JASP-aged 2 solely adopts the open D-loop state, a smaller fraction of F-actin-JASP-aged 1 also populates the open D-loop state (extra density protruding from Q41; also see Movie S1D-E; only the open conformation was modelled).

**Figure 5.**
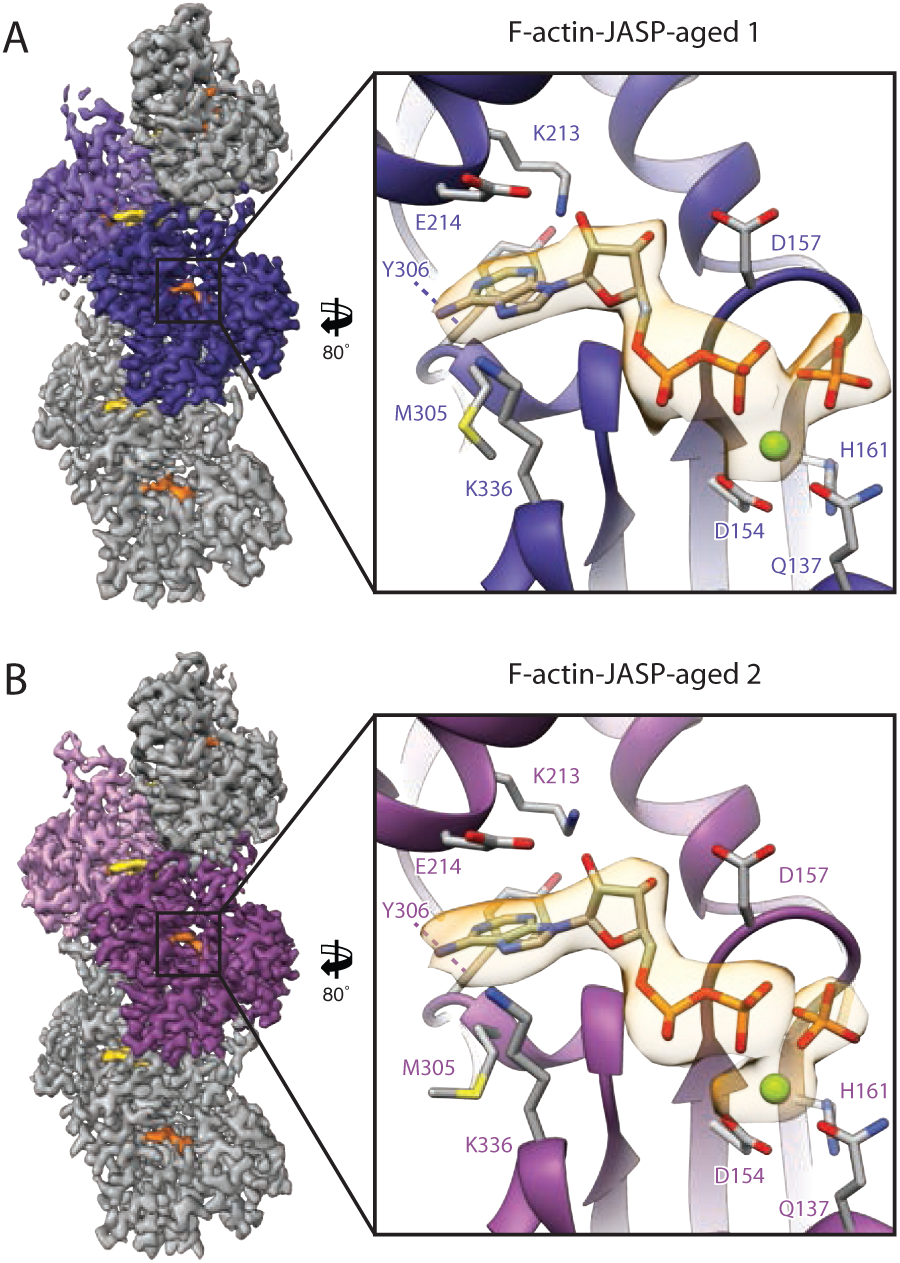
Nucleotides bound to JASP-stabilized F-actin. **(A-C)** Depiction of the central nucleotide binding cleft of F-actin-JASP-aged. In contrast to the phalloidin-stabilized F-actin structures, both F-actin-JASP-aged structures have strong density for an inorganic phosphate (P_i_) (also see Figure S2D-E and Movie S2D-E). Nucleotides and JASP are colored in orange and yellow, respectively. For color-code see Figure 4.

To verify our results and to exclude the possibility of a P_i_ contamination of the used glassware, we repeated the reconstitution of F-actin-JASP-aged, taking extra measures to avoid impurities. The resulting cryo-EM structure, which is resolved to an average resolution of ∼ 3.1 Å (Figure S1, Table 2), has the same strong density for P_i_ (Figure 5B, Movie S2I-J). The position of the inorganic phosphate and the organization of the active site are the same in both of the F-actin-JASP-aged data sets and resemble the one of F-actin-ADP-P_i_ (Figure S2D-E). The D-loop solely adopts the open conformation (Figure 4B, Movie S1I-J). We can therefore exclude that P_i_ impurities caused the strong P_i_ binding.

Taken together, the structure of F-actin-JASP-aged resembles our previous F-actin-ADP-P_i_-JASP structure (Merino et al., 2018) and differs considerably from that of F-actin-PHD-aged. We conclude that phalloidin and JASP do not share the same mode of action. Although we cannot exclude a more complex mechanism, it is most likely that JASP, in contrast to phalloidin, stabilizes the open D-loop state irrespective of the bound nucleotide or when it was added. The higher P_i_ affinity in the case JASP might account for the stronger stabilizing effect of JASP on F-actin in comparison to phalloidin (Visegrády et al., 2005; 2004).

### Comparison of phalloidin and JASP binding to F-actin

The structure and binding site of JASP is the same in all of our different cryo-EM structures of F-actin complexed with JASP (Merino et al., 2018; Pospich et al., 2017) (Figure 6, Movie S3G-J). Its binding site overlaps to a large proportion with the one of phalloidin, as illustrated by a superposition of structures (Figure S3D-F). The macro cycles of both molecules align well and stack onto the same hydrophobic patches on the actin surface. Furthermore, both molecules contain an indole group that occupies the same hydrophobic pocket. This indole group has been proposed to be of great importance for phalloidin binding (Mentes et al., 2018) and was shown to be essential for its toxicity (Falcigno et al., 2001; T. Wieland, 1977).

**Figure 6.**
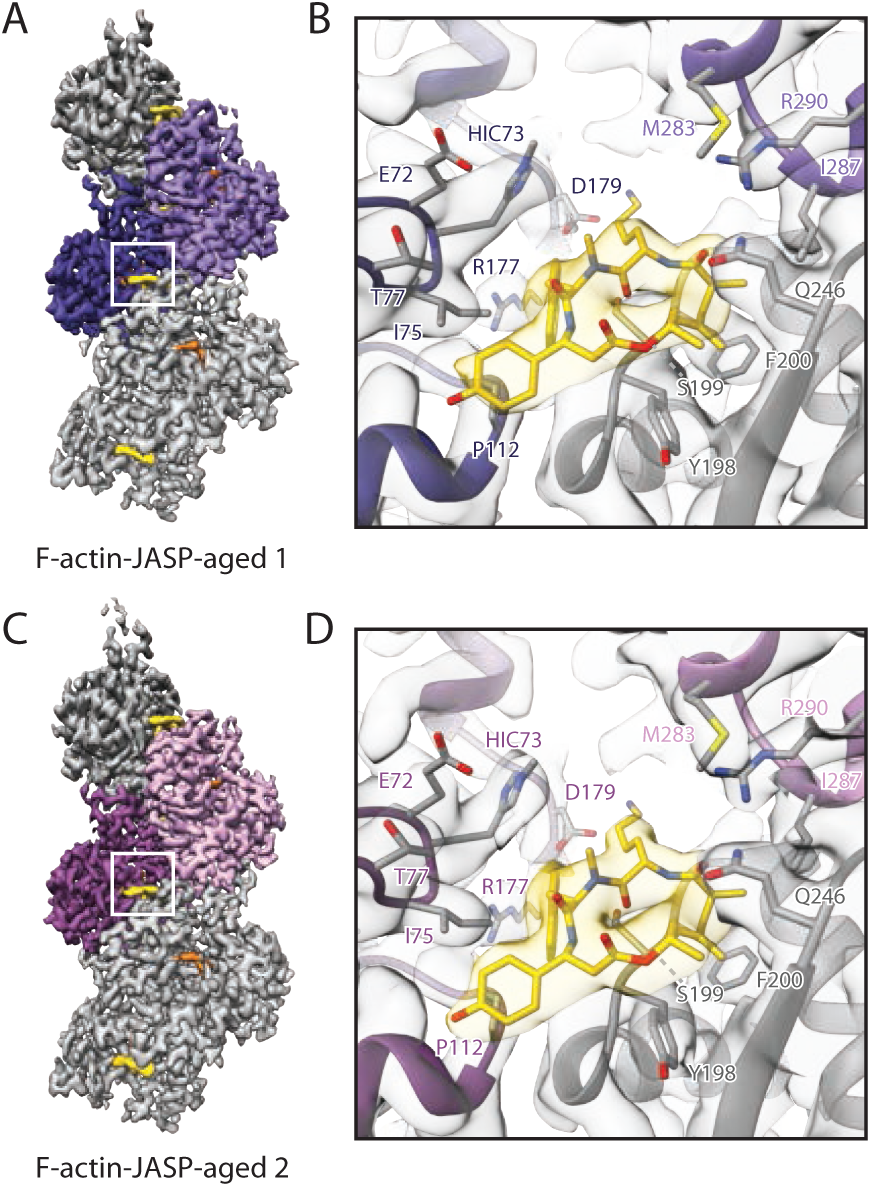
The Jasplakinolide binding site. Central binding sites illustrating the interaction of JASP (yellow) with three subunits from both strands. Phalloidin and JASP share the same binding site as highlighted in Figure S3D-F; also see Movie S3D-E, for color-code see Figure 4.

We have recently proposed that JASP inhibits P_i_ release by directly interacting with loops involved in phosphate release (Kabsch et al., 1990; Schulten and Wriggers, 1999) and by pushing SD1 toward the pointed end resulting in a stabilization of the open D-loop state (Merino et al., 2018). Despite the similarity of interactions of phalloidin and JASP with F-actin, we observe differences in the D-loop conformation between F-actin-PHD and F-actin-PHD-aged. Thus, the binding of phalloidin alone does not result in the open D-loop state by pushing SD1. Therefore, we propose that the interactions of phalloidin with loops near the active site are sufficient to explain the effects of the toxins.

### The structural effect of a fluorophore-conjugated phalloidin on F-actin

Fluorophore-conjugated phalloidin is widely used for labeling F-actin in biochemical and cellular experiments (Melak et al., 2017). To explore its structural effect on actin filaments, we solved the cryo-EM structure of F-actin copolymerized with Alexa Fluor 546 phalloidin (F-actin-PHD-Alexa) at ∼ 3.6 Å (Figure S1, Table 1). The structure reveals that Alexa Fluor 546 phalloidin binds to the same binding pocket as phalloidin with the fluorophore protruding from the filament (Figure 3G-I, Movie S3E-F). The Alexa 546 fluorophore is highly flexible and hence not resolved to a high resolution (Figure 3G). Its presence, however, becomes obvious when examining the map filtered to lower resolution (Figure 3I). Similar to F-actin-PHD, the active site contains ADP and at a lower occupancy P_i_ (Figure 2C, Movie S2C, Figure S2C). In contrast to F-actin-PHD, the density of the tip of the D-loop and the terminal part of the C-terminus are fragmented, suggesting a mixture of conformational states (Figure 1C, Movie S1E-F). The stem of the D-loop, however, indicates a higher population of the closed state in contrast to the open D-loop state in F-actin-PHD. We therefore believe that the conjugated fluorophore results in slower binding of phalloidin to F-actin, allowing for the aging of a subset of actin subunits before Alexa Fluor 546 phalloidin can bind.

### Conformational changes upon P_i_ release

The cryo-EM structures of F-actin-PHD and F-actin-PHD-aged represent the only pair of maps available where the open and the closed D-loop states are clearly linked to P_i_ binding. To identify conformational changes upon P_i_ release at the nucleotide binding site and how they are interconnected to the structural reorganization at the periphery of the filament, we compared the two structures. However, since the resolution of the available F-actin reconstructions is limited to ∼ 3 - 4 Å (Pospich and Raunser, 2018), subtle differences are often not reflected in the corresponding atomic models. We therefore carried out a thorough comparison of the two density maps (Movie S4).

In absence of P_i_, Mg^2+^ moves closer to the position previously occupied by P_i_. Moreover, both the phosphate binding loop 2 (residues 156-159) and the switch loop (residues 70-78) move towards the nucleotide, thereby partially closing the empty P_i_ site (Movie S4A-D). Orchestrated with the other loops, phosphate binding loop 1 (residues 14-16) and the proline-rich loop (residues 108-112) also move slightly closer to the nucleotide (Movie S4C,E). The importance of both phosphate binding loops as well as the switch loop was already proposed early on based on crystal structures, mutations in yeast and molecular dynamics (MD) simulations of G-actin (Belmont et al., 1999b; Kabsch et al., 1990; Otterbein et al., 2001; Schulten and Wriggers, 1999; Wriggers and Schulten, 1997). The conformational change of the loops at the active site is coupled to the surrounding subdomains and eventually results in a considerable downward movement of SD1 and SD2 (Movie S4F-H). This movement in turn leads to a major conformational change of the D-loop and C-terminus (changing from open to closed D-loop state) (Movie S4F-H). This is in line with several previous studies, suggesting that P_i_ release sets in motion a sequence of events resulting in a change of SD2 and structural rearrangement of the D-loop and C-terminus (Chou and Pollard, 2019; Graceffa and Dominguez, 2003; Otterbein et al., 2001; Saunders et al., 2014; Schulten and Wriggers, 1999; Zheng et al., 2007).

The organization of the D-loop C-terminus interface of F-actin (Ecken et al., 2015) suggests a cooperative transmission of this conformational change across neighboring subunits within one strand. In line with this, we observe a movement of SD3 that partially follows SD1 (Movie S4H). SD3 is linked to the SD2 of the next subunit via the W-loop (residues 165-172), which is thought to act as a nucleotide sensor (Kudryashov et al., 2010), primarily by Tyr169 inserting into the D-loop by a lock-and-key interaction (Ecken et al., 2015).

Arg177 has been proposed to directly bind to P_i_ and shuttle it out of the active site, involving a large displacement of the residue (Schulten and Wriggers, 1999). However, Arg177 is in the same position in both of our structures. This might be due to the binding of phalloidin, which is in close vicinity to Arg177. When mutated together with Arg179 to alanine, phalloidin does not bind to yeast F-actin anymore (Drubin et al., 1993), indicating that this residue is directly involved in phalloidin binding. In addition, phalloidin is in direct contact with both the switch and proline-rich loop (Figure 3, Movie S3A-F and Movie S4). Taken together, we therefore believe that the interaction of phalloidin with these specific loops and residues is the structural basis of the significant reduction of P_i_ release.

### States in stabilized filaments are representative for non-stabilized F-actin

Rather than inducing artificial states, phalloidin and JASP probably trap conformations that occur during the natural aging process of F-actin. Thus, we believe that the structural changes that we observed in phalloidin-stabilized F-actin represent those in non-stabilized F-actin. To test this hypothesis, we performed a principle component (PC) analysis of atomic models and correlated structures of toxin-stabilized F-actin to structures of F-actin in different nucleotide states (Figure 7). The resulting distribution in the PC space indicates a close similarity of structural states and allows one to classify the conformational differences.

**Figure 7.**
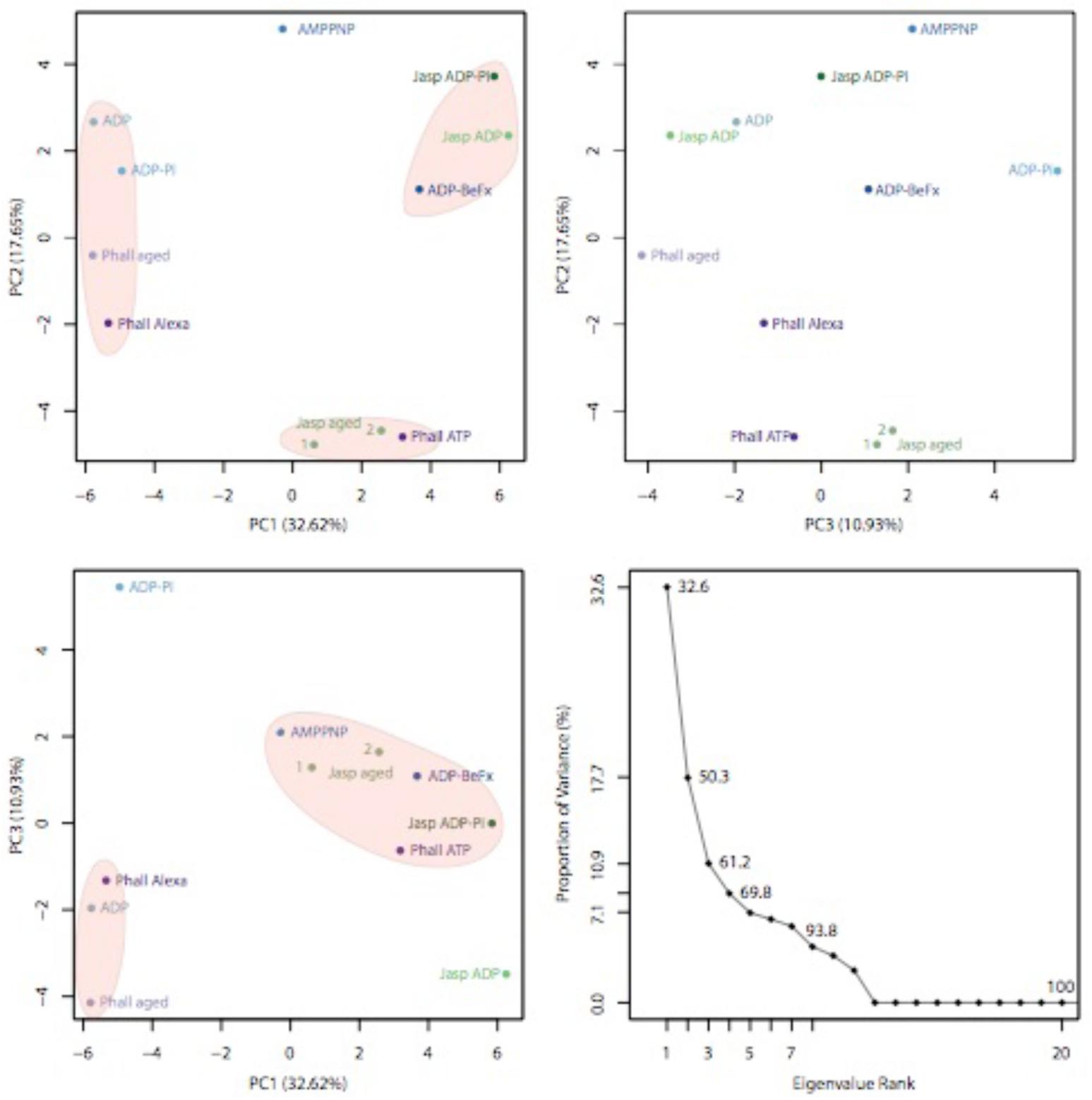
Principle component analysis of atomic models of F-actin. Principle component analysis including phalloidin (shades of purple) and JASP-stabilized (light green) F-actin structures presented in this manuscript, F-actin bound to different nucleotide states, namely AMPPNP, ADP-BeFx, ADP-Pi and ADP (shades of blue) (Merino et al., 2018) and filaments stabilized by JASP, including JASP ADP-Pi and JASP ADP (shades of green) (Merino et al., 2018) – PDB accession numbers listed in methods. The percentage of the total variance attributed to each component is shown in the axis labels and summarized in the panel at the right bottom. Prominent clusters are highlighted by orange ellipsoids. Data points are labeled with abbreviations based on the bound nucleotide and/or stabilizing agent. Also see Movies S5-S7.

Models are arranged according to their D-loop state along the first PC, with the closed D-loop state at one end and the open D-loop state at the other. In line with this, the trajectory along the first PC (Movie S5) illustrates an upwards rotation of SD1 and SD2 and the reorganization of the D-loop and C-terminus, similar to what we have previously reported (Merino et al., 2018). The arrangement of atomic models along the second PC is less clear and does not display clearly separated clusters. The trajectory along this component (Movie S6) displays a rotation or torsion of SD1 and SD2 against SD3 and SD4, that we currently cannot connect to any characteristic of the mapped structures. In case of the third PC, models are arranged according to the occupancy of the inorganic phosphate site in the nucleotide binding pocket. Models with only ADP are located at one end, while F-actin-ADP-P_i_, which likely has the highest P_i_ occupancy, marks the other end. The trajectory along the third PC illustrates a breathing motion of the complete protomer, where the release of P_i_ results in a more compact structure (Movie S7).

The structural changes along the first and third PC reflect the rearrangements that we identified for phalloidin-stabilized filaments upon P_i_ release. The relative positions of our phalloidin F-actin structures - close to the ones complexed with ADP-BeF_x_ and ADP, respectively - support our hypothesis that phalloidin does not induce artificial conformations but instead stabilizes states within the natural conformational space of F-actin (Figure 7). Thus, our findings with toxin-stabilized F-actin are probably representative for actin filaments in general.

## Conclusions

In this study we demonstrate that the stabilizing agent phalloidin interferes with the natural aging process of F-actin, which is key to its cellular functions (Figure 8). Phalloidin traps different conformational states depending on when it is added to actin. Addition of phalloidin to actin immediately before polymerization results in the stabilization of the open D-loop state and the reduction of P_i_ release. However, when phalloidin is added to aged filaments that have already released P_i_, it stabilizes the closed D-loop state. Thus, phalloidin cannot revert the conformational change associated with P_i_ release once it has taken place. While we found that the Alexa 546 fluorophore alters the effect of phalloidin on F-actin, it still affects the aging process of the filament. JASP also prevents F-actin from aging by trapping it in a young state and inhibiting P_i_ release, but it does not act precisely in the same way as phalloidin. Notably, it dramatically increases the affinity of F-actin for P_i_ (Figure8). While cryo-EM structures are well suited to illustrate the effects of stabilizing toxins on the natural aging process of F-actin, they fail to shed light onto the underlying dynamic processes. Thus, to fully understand the mode of action of phalloidin and JASP, future TIRF microscopy experiments or binding assays similar to what was done for coronin, myosin 5 and myosin 6 (Cai et al., 2007; Merino et al., 2018; Zimmermann et al., 2015) should be conducted.

**Figure 8.**
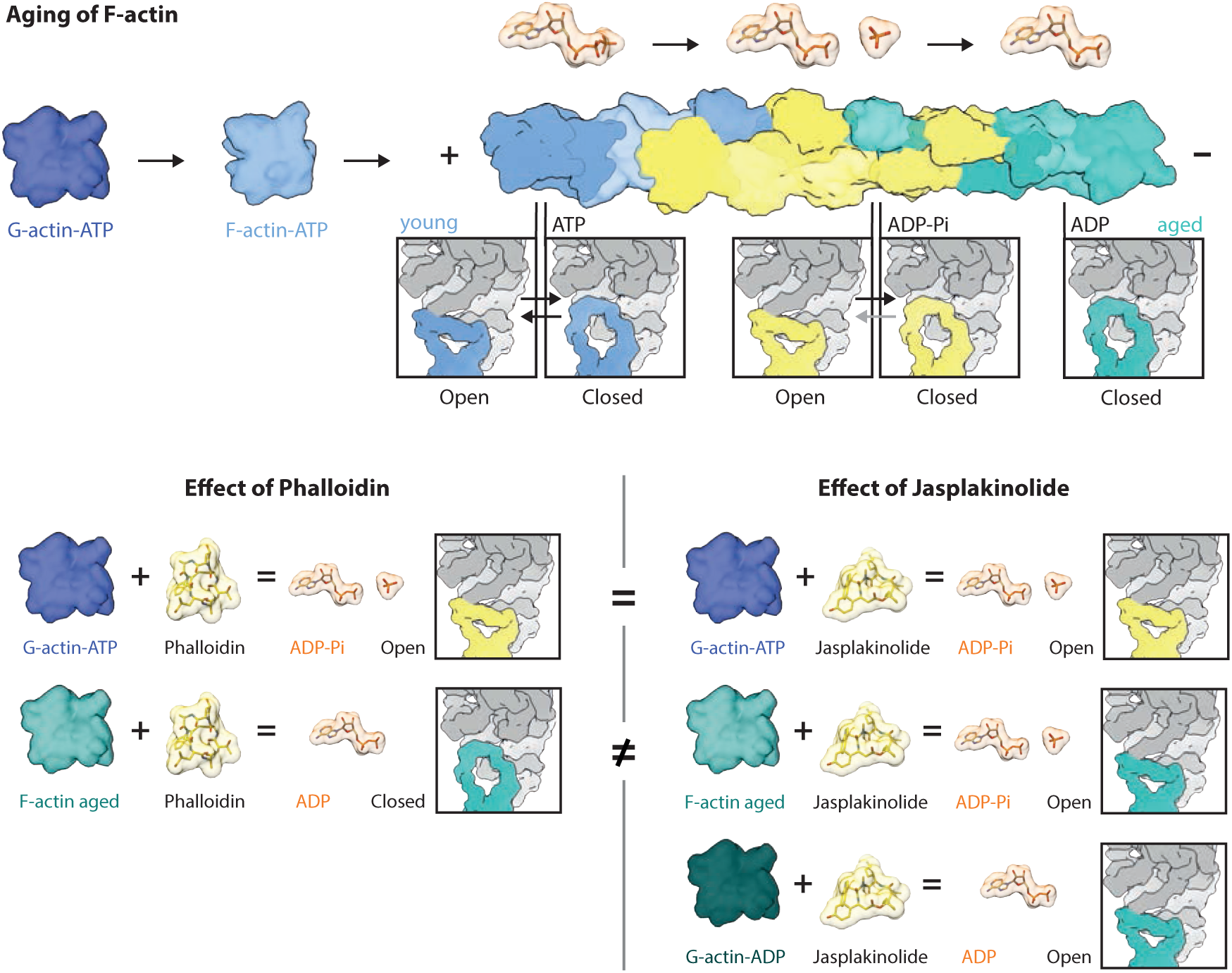
Schematic illustration of the effect of phalloidin and JASP on F-actin. (Top Panel) Summary of actin polymerization and filament aging highlighting associated structural states and their dependence on the bound nucleotide. **(Bottom Panel)** Depiction of the effect of phalloidin **(left)** and JASP **(right)** on F-actin as elucidated by cryo-EM reconstructions. While phalloidin and JASP have the same effect when added to G-actin-ATP before polymerization, their effect significantly differs when added after aging of filaments.

Our results have strong implications on the usage of both phalloidin and JASP for the stabilization and labeling of actin filaments in electron or fluorescent light microscopy. While it is known that both toxins affect actin dynamics, we demonstrate that binding of phalloidin and JASP interferes with the natural aging process of F-actin. This likely alters the interactions with ABPs, especially interactions with partners that sense the nucleotide state of actin to enable directed transport or remodeling of the cytoskeleton (Cai et al., 2007; Ge et al., 2014; Suarez et al., 2011; Zimmermann et al., 2015). While we do not know if some ABPs might be able to overwrite the effect of phalloidin and JASP, we believe that the toxins disrupt the balance within the complex actin cytoskeletal system.

Phalloidin is commonly used at substoichiometric concentrations for fluorescent labeling, thereby its full negative impact is possibly concealed. It is, however, difficult to assess the extent of the effect of phalloidin in a given experiment, especially when considering the proposed cooperative effect of phalloidin (Orlova et al., 1995; Visegrády et al., 2005). Because of the described possible disadvantages of phalloidin and JASP, we suggest that alternative labels are reconsidered, especially when investigating actin dynamics and protein interactions. There are many different labeling techniques available; based on either small molecules, peptides from ABPs, anti-actin-nanobodies or fusion constructs to a fluorescent protein or tag (for reviews see (Belin et al., 2014; Melak et al., 2017)).

Furthermore, our work illustrates that the stabilization of filaments likely does not result in artificial states but traps intermediate ones within the natural conformational space of F-actin. Thus, we believe that toxin-stabilized F-actin can be used to mimic a certain nucleotide state and that structural rearrangements observed in these filaments are representative for conformational changes of F-actin in general.

## Acknowledgements

We thank O. Hofnagel and D. Prumbaum for assistance with data collection. We thank W. Linke and A. Unger (Ruhr-Universität Bochum, Germany) for providing us with muscle acetone powder. We thank H.-D. Arndt for providing us with jasplakinolide (Jasp-cLys). This work was supported by the Max Planck Society (to S.R.) and the European Council under the European Union’s Seventh Framework Programme (FP7/ 2007–2013) (grant 615984) (to S.R.). S.P. was supported as a fellow of Studienstiftung des deutschen Volkes.

## Author contributions

Conceptualization: S.P. and S.R. Investigation: S.P. and F.M. Formal analysis: S.P. Validation: S.P. and F.M. Visualization: S.P. Writing – Original Draft: S.P. Writing – Review & Editing: S.P., F.M. and S.R. Supervision and funding acquisition: S.R.

## Materials and Methods

### Protein purification and sample preparation

Actin was purified from rabbit skeletal muscle acetone powder using several polymerization and depolymerization steps as described previously (Merino et al., 2018; Pardee and Spudich, 1982). Purified G-actin was stored in G-actin buffer (5 mM Tris pH 7.5, 1mM DTT, 0.2 mM CaCl_2_, 2mM NaN_3_ and 0.5 mM ATP) at −80 °C. Aliquots were freshly thawed and cleared through centrifugation, before polymerization was induced by adding 100 mM KCl, 2mM MgCl_2_ and 0.5 mM ATP. An additional cycle of depolymerization/polymerization was performed for F-actin-PHD and F-actin-PHD-Alexa samples. Phalloidin (Sigma Aldrich), phalloidin Alexa Flour 546 (Thermo Fischer) and jasplakinolide (JASP, see (Merino et al., 2018) for details) were solved in methanol and DMSO, respectively, producing a 2-3 mM stock solution. For F-actin-PHD and F-actin-PHD-Alexa samples a ∼ 2-fold molar excess of ligand was added directly before induction of the polymerization. After overnight polymerization filaments were collected through ultracentrifugation and pellets were rinsed and resuspended in F-buffer (5 mM Tris pH 7.5, 1 mM DTT, 100 mM KCl, 2mM MgCl_2_ and 2mM NaN_3_) with additional small amounts of phalloidin and phalloidin Alexa Flour 546, respectively, to avoid falling below K_D_. Aged samples (F-actin-PHD-aged, F-actin-JASP-aged) were polymerized overnight in absence of any stabilizing small molecule to ensure complete hydrolysis of ATP. Finally, phalloidin or JASP were added in ∼ 2-fold molar excess after multiple rounds of careful rinsing (with F-actin buffer to reduce amount of P_i_) and resuspension of the pellet in F-actin buffer. Note that the two F-actin-JASP-aged data sets are replicates that were prepared independently. Additional 0.2 mM ADP and 0.02 v/w% Tween 20 were added to F-actin-JASP-aged samples.

### Cryo-EM grid preparation and screening

An initial sample check and adjustment of the protein concentration was performed for all samples except F-actin-PHD-aged and F-actin-JASP-aged 2 using a standard negative staining protocol described before (Ecken et al., 2015). Screening was performed on a Tecnai G Spirit microscope (FEI Thermo Fisher) operated at 120 kV and equipped with a LaB_6_ cathode and a CMOS TemCam F416 (TVIPS) camera. After initial screening of the F-actin-PHD sample, filaments were spun down and stored as a pellet at 4°C for ∼ 2 weeks. After resuspension and addition of supplementary phalloidin to ensure saturation, filaments were screened again and found to be indistinguishable from those seen in the first screening.

For cryo grid preparation 0.02-0.03 v/w% of Tween 20 were added to all samples if not yet done before. All samples were diluted in F-buffer without additional nucleotide or stabilizing small molecule resulting in the following final protein and methanol/DMSO concentrations; F-actin-PHD ∼ 0.2 µM, 0.01%, F-actin-PHD-Alexa ∼ 8.4 µM, 1.48%, F-actin-PHD-aged ∼ 5 µM, 2.0%, F-actin-JASP-aged 1 ∼ 7.5 µM, 0.5 % and F-actin-JASP-aged 2 ∼ 5 µM, 0.2 %. Values are estimates based on the initial, spectrophotometrically determined concentrations.

In case of F-actin-PHD, 1.5 µl sample were incubated for 30 s on a previously glow discharged holey carbon grid (QF R2/1 300 mesh, Quantifoil) and manually backside blotted for 9s using Whatman no. 5 filter paper. Grids were then directly plunged into liquid ethane using a CP3 cryo plunger (Gatan) at ∼ 98 % humidity. All other samples were blotted and plunged automatically using a Vitrobot cryo plunger (FEI Thermo Fischer) operated at 13 °C and 100% humidity. 1.5 - 3 µl of sample were applied onto a previously glow discharged holey carbon grid (QF R2/1 300 mesh, Quantifoil) and directly blotted for 6.5 - 9.0 s using a blotting force of −20 or −25. After a drain time of 0 - 1s grids were automatically plunge frozen into liquid ethane. Prescreening and cryo sample optimization was performed either on a Tecnai G Spirit microscope (FEI Thermo Fisher) operated at 120 kV using a single-tilt cryotransfer holder 626 (Gatan) or on a Talos Arctica (FEI Thermo Fisher) operated at 200 kV and equipped with a Falcon III direct detector (FEI Thermo Fisher).

### Cryo-EM data acquisition

Data sets were collected on Titan Krios microscopes (FEI Thermo Fisher) operated at 300 kV and equipped with a X-FEG using EPU. Equally dosed frames were collected using Falcon II, Falcon III (linear mode, FEI Thermo Fisher) or K2 Summit (super-resolution mode, Gatan) direct electron detectors, the latter in combination with a GIF quantum-energy filter set to a filter width of 15 eV. For every hole four images with carbon edge were collected. Acquisition details of all five data sets including pixel size, electron dose, exposure time, number of frames and defocus range are summarized in Table 1 and Table 2. Data collection was monitored live using TranSphire (Stabrin, 2018), allowing for direct adjustments of data acquisition settings when necessary i.e. defocus range or astigmatism. The total number of images collected is summarized in in Table 1 and Table 2.

### Image processing of cryo-EM data sets

Collected movies were automatically and on-the-fly preprocessed using TranSphire (Stabrin, 2018). Preprocessing included drift correction with MotionCor2 (Zheng et al., 2017) creating aligned full-dose and dose-weighted micrographs. In this step the JASP aged 2 super-resolution images were furthermore binned twice. For all other data sets additional aligned micrographs with a total dose of ∼ 25 e/Å^2^ and ∼15 e/Å^2^ were generated. CTF estimation was also performed within TranSphire; running either CTFFIND4.1.5 or GCTF (only for F-actin-PHD-Alexa and F-actin-JASP-aged 2) on non-dose weighted aligned micrographs. Unaligned frame averages were manually inspected and judged in terms of ice and protein quality, resulting in a removal of 19-30 % of the data sets, see Table 1 and Table 2 for details.

Particles (F-actin-PHD and 80% of F-actin-PHD-Alexa) were selected manually on bandpass filtered images with a box size of 256 px and a particle distance of 37 px using sxhelixboxer in SPARX (Hohn et al., 2007). Filaments were only included when they contained at least six segments. Remaining images of the F-actin-PHD-Alexa data set, F-actin-PHD-aged and F-actin-JASP-aged 1 were auto-picked using STRIPPER (Wagner, 2018) with a box size of 256 px and a particle distance of 40 px and 32 px, respectively. The F-actin-JASP-aged 2 data set was auto-picked using the filament mode of crYOLO (Wagner et al., 2019) with a filament width of 60 px, box distance of 27 px, a minimum number of six segments and on-the-fly lowpass filtering. To train the crYOLO model 94 micrographs were initially picked manually using sxhelixboxer resulting in ∼ 15k particles.

All following processing steps were performed in Relion-2-beta or Relion-3 (JASP aged 2) (Scheres, 2012; Zivanov et al., 2018). Particles were extracted from dose weighted, aligned micrographs for all data sets using a box size of 256 px. For F-actin-PHD and F-actin-JASP-aged 1 particles were additionally binned to 192 px resulting in a pixel size of 1.52 Å/px and 1.5 Å/px, respectively. For all data sets except F-actin-JASP-aged 2, particles were furthermore extracted from micrographs with a total dose of 25 e/Å^2^ and 15 e/Å^2^ using a box size of 256 px. The total number of extracted particles is listed in Table 1 and Table 2.

For auto-picked particles (subset of F-actin-PHD-Alexa containing 194,777 particles, F-actin-PHD-aged and both data sets of F-actin-JASP-aged) helical 2D classification was used to discard false picks and particles of bad quality. As removal of particles can result in gaps within filaments and this would cause problems during prior calculations (see below), star files were cleaned up by either splitting filaments into two new filaments or removal if split filaments would contain less than six segments. The remaining number of particles was 184,566 for the F-actin-PHD-Alexa subset (merged data set contains 895,735 particles), 1,078,848 for F-actin-PHD-aged, 768,490 for F-actin-JASP-aged 1 and 415,892 for F-actin-JASP-aged 2. Manually picked particles were not 2D classified.

To speed the refinement up, the number of particles for F-actin-PHD-aged and F-actin-JASP-aged 1 was further reduced by removing all micrographs with a CTF maximum resolution worse than 4.5 Å and 6.5 Å, respectively, a total drift higher than 15 Å or a per frame drift higher than 1.12 Å (1 px) resulting in a total of 646,699 and 360,799 particles, respectively.

As helical refinement tends to result in over refinement and loss of detail due to averaging, we used the modified single-particle 3D refinement approach we described in detail before (Merino et al., 2018). To enable single particle refinement helical priors and flip ratios were removed from the star files and helical half sets introduced manually. An initial 3D reference was created from PDB 5JLF (after removal of tropomyosin) (Ecken et al., 2015; 2016) and filtered to 25 Å using SPARX (Hohn et al., 2007) and EMAN2 (Tang et al., 2007). The initial refinement was performed on binned or full-size dose-weighted particles with a sampling of 3.7° limiting the tilt angle to 89° (--limit_tilt 89). As the sampling rate was not automatically increased to 1.8° for the F-actin-PHD-Alexa and F-actin-PHD-aged data sets, a follow up refinement using a sampling of 1.8 ° and the previous output map filtered to 7-8 Å as reference was carried out. Output maps were afterwards filtered to 6-7 Å and used as reference for a local refinement with a sampling of 0.9° without any angular restraints. Based on those projection parameters, prior values for the tilt and in plane rotation angles were calculated and outliers discarded (see (Merino et al., 2018) for details), resulting in a total of 529,081 (F-actin-PHD), 719,053 (F-actin-PHD-Alexa), 513,783 (F-actin-PHD-aged), 352,285 (F-actin-JASP-aged 1) and 336,783 (F-actin-JASP-aged 2) particles.

All following processing steps were performed on unbinned particles with a total electron dose of 25 e/Å^2^. Reference maps and star files were rescaled and modified accordingly. Previously calculated prior values were used as prior distribution restraints (--sigma_tilt 1 and –sigma_psi 1) in the subsequent refinement using a sampling of 0.9° and reducing the shift range. Scaled maps filtered to 4-5 Å were used as references. Finally, the map quality was further improved by continuing the last iteration using particles with a total dose of 15 e/Å^2^. In case of the F-actin-JASP-aged 2 data set the electron dose was not reduced as described before. Instead CTF refinement and bayesian polishing was performed in Relion-3 (Zivanov et al., 2018). Polished particles were afterwards used for the final local refinement with prior restraints.

All refinements were performed using a mask. Initial refinements made use of a mask spanning 85% percent of the filament with an extension of 3 px and softedge of 5 px. Afterwards a mask covering a central sphere with a radius of ∼ 110 px, an extension of 4 px and softedge of 10 px was used in combination with solvent flattening of the FSC.

In case of F-actin-PHD, the map quality could be further improved by removing all micrographs having an overall drift above 6 Å and running a 2D classification without alignment to remove all classes without features. The last iteration was finally repeated with this subset containing a total of 405,265 particles. In case of F-actin-PHD-Alexa, the refinement seemed not to have converged yet. Thus, the final map could be improved by another local auto refinement using the same particle subset and settings as used for the previous final iteration.

For the final post processing we generated a mask containing the central ∼120 Å of the filament (as done before (Merino et al., 2018)) resulting in 3.31 Å for F-actin-PHD, 3.58 Å for F-actin-PHD-Alexa, 3.67 Å for F-actin-PHD-aged, 3.68 Å for F-actin-JASP-aged 1 and 3.09 Å for F-actin-JASP-aged 2. Final resolutions, number of micrographs before and after sorting as well as particles numbers are summarized in Table 1 and Table 2.

### Local resolution estimation and filtering

Local resolutions were calculated on half maps using Sphire (Moriya et al., 2017), providing the final average resolution reported by Relion. Half maps sharpened to the automatically determined B-Factor in Relion’s postprocessing were afterwards filtered to local resolution in Sphire.

### Model building and refinement

An initial atomic model and cifs library for phalloidin was generated using elBow (Moriarty et al., 2009) in Phenix (Adams et al., 2011) inputting the SMILES string provided by the ZINC data base (Irwin and Shoichet, 2005). This model was then manually fitted using cifs restraints into the central density of phalloidin within Coot (Emsley et al., 2010). The orientation was unambiguous and in good agreement with a recently published model of an actomyosin complex including phalloidin (Mentes et al., 2018). Initial F-actin models for F-actin-PHD and F-actin-PHD-Alexa are based on PDB 5OOD (F-actin-ADP-Pi, (Merino et al., 2018)). As the occupancy of P_i_ is low it was removed and Jasp-cLys replaced by the previously generated initial model of phalloidin. PDB 5ONV (F-actin-ADP (Merino et al., 2018)) was used as a starting model for F-actin-PHD-aged and phalloidin added manually. The model of F-actin-JASP-aged 2 is based on the final model of F-actin-PHD, where phalloidin was replaced by JASP and P_i_ added, both from PDB 5OOD. Models were docked into the density and adjusted where necessary using Chimera (Pettersen et al., 2004) and Coot (Emsley et al., 2010). Finally, the central five subunits were fitted individually into the density, HIC73 replaced by HIS for phalloidin structures and hydrogens added. In all following steps five subunits were included.

Consecutive model building and refinement was performed in Rosetta (Wang et al., 2016). To avoid overfitting, we modelled against one half map (sharpened, broadly masked and filtered to resolution) and used the other for validation. Helical symmetry was enforced in all following steps and additional distance constraints applied to ensure proper modelling of Mg^2+^ and P_i_. For ligands, Rosetta creates an approximate parameter set from the molecule’s chemical nature, assuming that the starting model is close to an energy minimum. For all models except F-actin-JASP-aged 1 we used between one and three rounds of iterative rebuilding, generating 960 models every round. After convergence assessment, the best scoring model was locally refined and relaxed using Rosetta with minimization and repacking. When necessary models were adjusted manually in Coot, followed by another round of iterative rebuilding or local refinement. Although F-actin-JASP-aged 1 adopts both the open and closed D-loop state, only the dominant open state was modeled. Considering that F-actin-JASP-aged 1 is a replicate of F-actin-JASP-aged 2, we used the final model of F-actin-JASP-aged 2 as a starting model and directly ran a local relaxation after minor adjustments in Coot. Final models were selected based on the best energy, FSC free, MolProbity (Chen et al., 2010) and EmRinger (Barad et al., 2015) scores (both calculated calling Phenix functions). Finally, we methylated HIS73 to HIC in all models of phalloidin-stabilized F-actin and regularized it in its ideal geometry in Coot. For F-actin-PHD-Alexa we removed residues 43-48 and 374-375 as the tip of the D-loop and the C-terminus are not sufficiently resolved. Bond and angle outliers of the F-actin-JASP-aged models were manually corrected in Coot. Finally, we utilized Rosetta for validation, to impose helical symmetry and calculate B-factors for the two modified F-actin-JASP-aged models. Refinement statistics of all five models are summarized in Table 1 and Table 2.

### Principle component analysis

Principle component analysis (PCA) was performed using the bio3D library (Grant et al., 2006) in R (R Core Team, 2017) using the RStudio Interface (RStudio Team, 2015). Models of F-actin actin in different nucleotide states – AMPPNP PDB: 5OOE, ADP-BeFx PDB:5OOF, ADP-Pi PDB: 6FHL and ADP PDB: 5ONV (Merino et al., 2018) and in complex with JASP – bound to ADP-Pi PDB: 5OOD or to ADP PDB: 5OOC (Merino et al., 2018) were included in addition to the models presented in this manuscript. Only the central actin chain of every model was used and ligands excluded. Models were aligned with the *pdbaln* method which uses the MUSCLE algorithm (Edgar, 2004) for sequence alignment. For superposition of models a stable core of residues was identified with *core*.*find* and afterwards used for superposition using *pdbfit*. Gaps in the sequence i.e. within the D-loop or C-terminus were excluded in the following PCA running *px*.*xray*. Data points were manually grouped for coloring based on bound proteins or ligands. Trajectories along the principle component axis were created using *mktrj*.*pca*. Note that gaps within the D-loop are not visible in the trajectory model as remaining atoms were renumbered building a continuous chain.

### Structure Visualization

The central subunit or central three subunits were used for the visualization of models and density maps, as they include all important contact sites and are best resolved. For all surface representations, models protonated with H++ (Anandakrishnan et al., 2012) at pH 7.5 with HIC replaced by HIS were displayed. When not indicated differently, maps filtered to local resolution where used for all figures. Figures and movies were created with in Chimera (Pettersen et al., 2004) and modified when necessary using image or movie processing software. Protein surfaces were colored by hydrophobicity using “define attribute” inputting amino acid-specific scores (Hessa et al., 2005).

### Data availability

The atomic models and cryo-EM maps are available in the PDB (Burley et al., 2018) and EMDB databases (Lawson et al., 2011), under accession numbers PDB XXX and EMD-XXXX (F-actin-PHD); PDB XX and EMD-XX (F-actin-PHD-aged PDB XXXX and EMD-XXXX (F-actin-PHD-Alexa); PDB XXXX and EMD-XXX (F-actin-JASP-aged 1); and PDB XXXX and EMD-XXX (F-actin-JASP-aged 2). The datasets generated during and/or analyzed during the current study are available from the corresponding author upon reasonable request.

## Supplementary information

**Figure S1.**
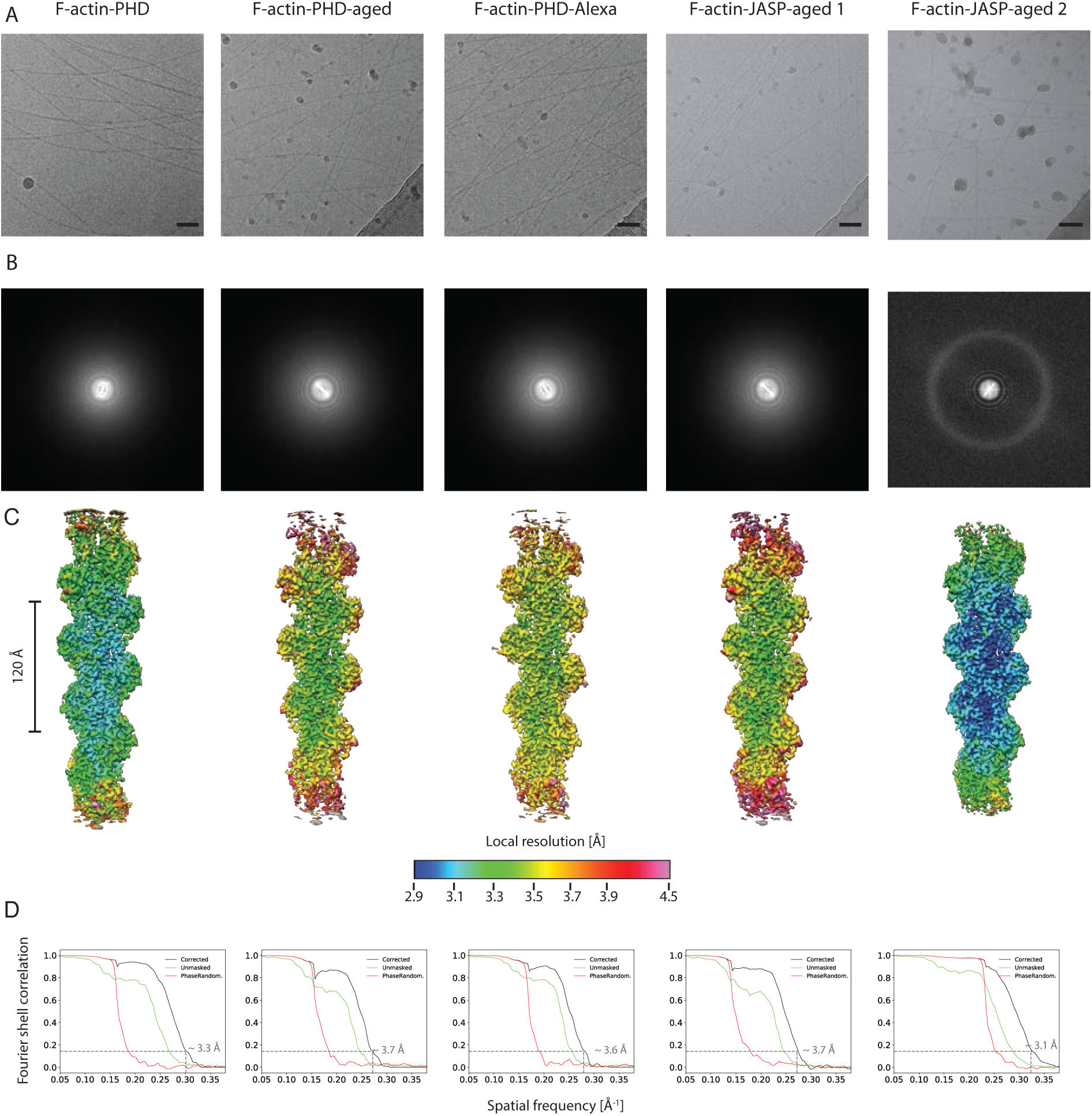
Overview of original cryo-EM data and resolutions. Related to Table 1 and Table 2. **(A**) Representative micrographs at ∼ −1.5 µm defocus (scale bar 500 Å) **(B)** and their power spectra. **(C)** Local resolution of all reconstructions illustrating the intrinsic flexibility of F-actin resulting in a gradual decrease of resolution towards the ends of the filament. **(D)** Fourier shell correlation (FSC) curves for the masked (black), unmasked (green) and phase-randomized (red) maps. While the FSC plots for the unmasked maps clearly show that there is no over-refinement in the reconstructions, the phase-randomized curves prove that the masking has no effect on the estimated resolution. FSCs were calculated for a 120-Å-long central section of the filament.

**Figure S2.**
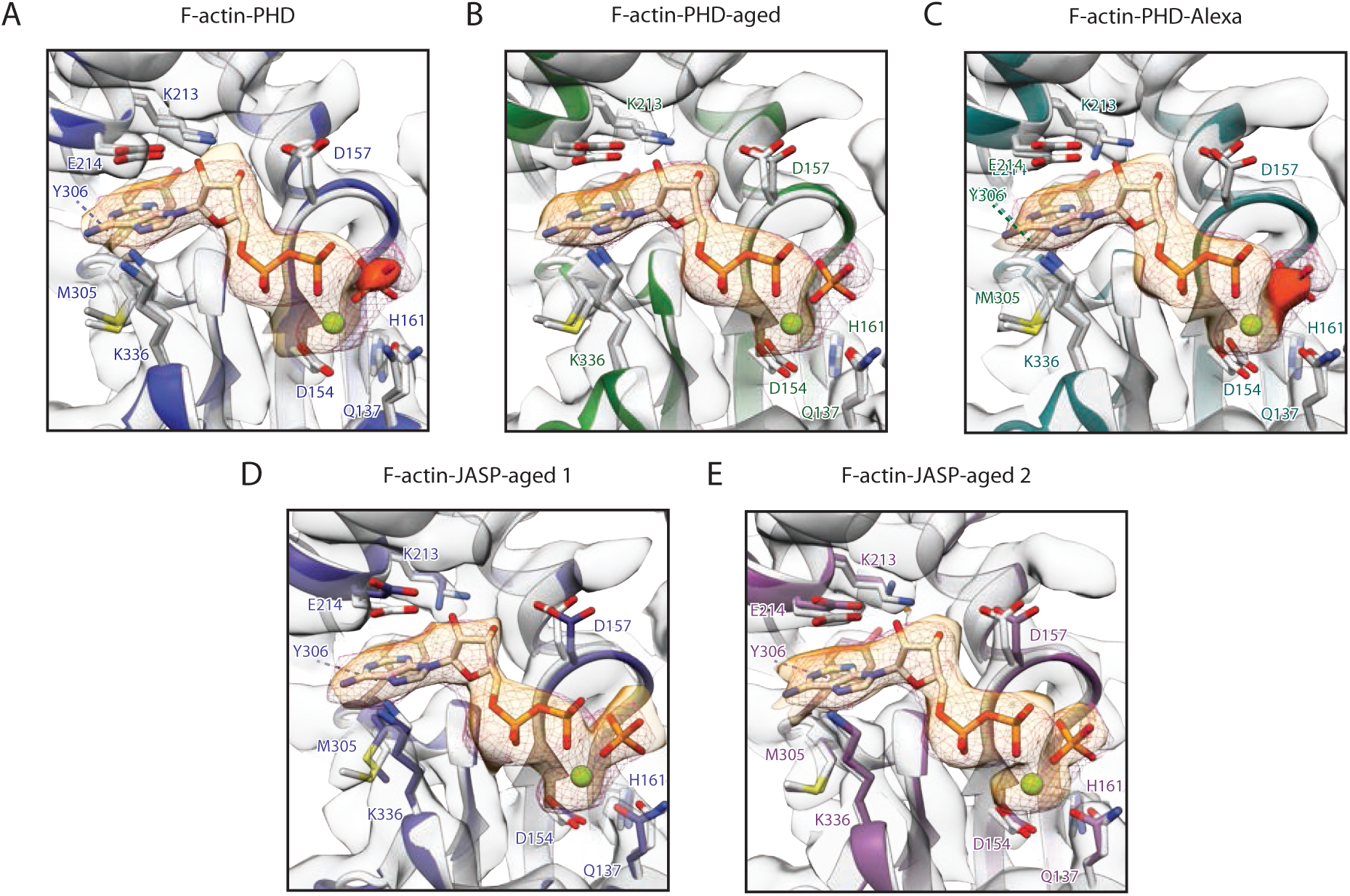
Comparison of nucleotide binding clefts. Related to Figure 2 and Figure 5. **(A-E)** Overlay of atomic models of **(A)** F-actin-PHD (blue), **(B)** F-actin-PHD-aged (green), **(C)** F-actin-PHD-Alexa (turquoise), **(D)** F-actin-JASP-aged 1 (purple) and **(E)** F-actin-JASP-aged 2 (lavender) with F-actin-ADP-P_i_ (grey, PDB: 6FHL (Merino et al., 2018)). For clarity only the atomic model of the nucleotide of F-actin-ADP-P_i_ is shown. The nucleotide density of F-actin-ADP-P_i_ (EMDB: 4259: (Merino et al., 2018)) is additionally shown as a pink mesh to highlight the absence (F-actin-PHD-aged) or consistency of the position of the inorganic phosphate (P_i_) (all other structures). As the density of P_i_ is weak for F-actin-PHD and F-actin-PHD-Alexa (highlighted by a darker shade of orange) it was not included into the corresponding atomic models. Superposition of models furthermore illustrates the conservation of the nucleotide binding pocket and active site. Shown densities (grey with the nucleotide highlighted in orange) are central subunits of locally filtered maps from this manuscript.

**Figure S3.**
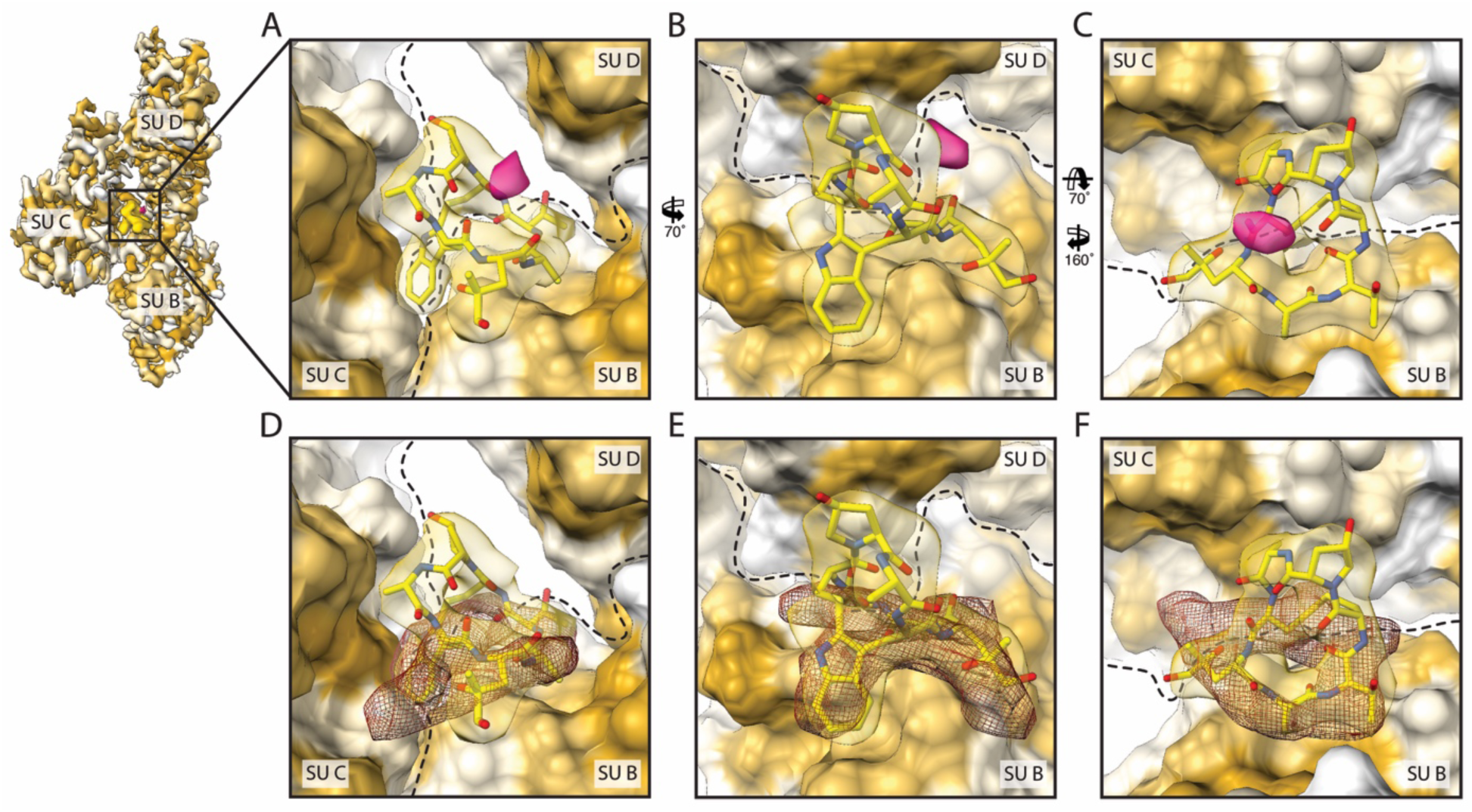
Hydrophobic nature and overlap of phalloidin and JASP binding sites. Related to Figure 3 and Figure 6. **(A-B)** Side and **(C)** tilted top views of the central phalloidin binding site highlighting large hydrophobic patches on all three subunits of F-actin. F-actin is depicted as surface and colored by hydrophobicity from high (yellow) to low (white, see Methods for details). Dashed lines indicate boundaries of actin subunits. The extra density, possibly corresponding to an ion is depicted in pink. **(D-F)** Same views as in **(A-C)** with an additional overlay of JASP (red mesh, EMDB: 3837 (Pospich et al., 2017)) illustrating that phalloidin (yellow) and JASP do not only share their binding site but also a key hydrophobic indole group. Also see Movie S3.

**Figure S4.**
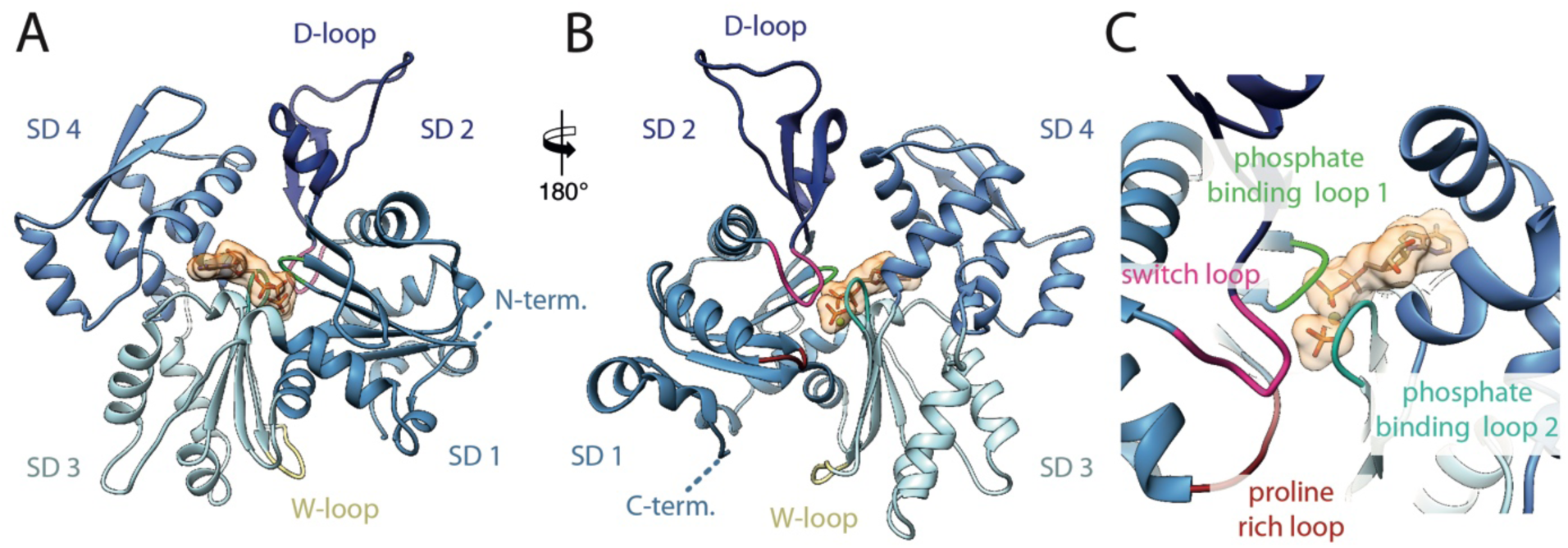
Overview of subdomain organization and localization loops associated with P_i_ release. Related to Movie S4. Front (**A**) and back (**B**) view of the F-actin protomer illustrating the domain organization and location of the termini and important loops. To highlight the nucleotide in the central cleft it is displayed as s surface (orange). (**C**) Close-up view of the backside of the nucleotide pocket with emphasize on loops in close vicinity.

## SI Movie legends

**Movie S1. Conformations at the intra-strand interface, Related to Figure 1 and Figure 4**.

Illustration of the intra-strand interface of all structures. The interface is made up by the D-loop and C-terminus of adjacent subunits within one strand and can adopt the open (F-actin-PHD, F-actin-JASP-aged 2) or closed state (F-actin-PHD-aged) as well as a mixture of both (F-actin-PHD-Alexa, F-actin-JASP-aged 2). Filament overview and close-up view of the interface of **(A-B)** F-actin-PHD (shades of blue), **(C-D)** F-actin-PHD-aged (shades of green), **(E-F)** F-actin-PHD-Alexa (shades of turquoise), **(G-H)** F-actin-JASP-aged 1 (shades of purple) and **(I-J)** F-actin-JASP-aged 2 (shades of lavender). Nucleotides and phalloidin/JASP are colored in orange and yellow, respectively.

**Movie S2. Nucleotides bound to phalloidin and JASP-stabilized F-actin structures, Related to Figure 2 and Figure 5**.

Illustration of the nucleotide binding cleft of all structures allowing the unambiguous identification of the bound nucleotide (orange) as either ADP (F-actin-PHD-aged) or ADP-P_i_ (all structures with different P_i_ occupancies). Filament overview and close-up view of the binding site of **(A-B)** F-actin-PHD (shades of blue), **(C-D)** F-actin-PHD-aged (shades of green), **(E-F)** F-actin-PHD-Alexa (shades of turquoise), **(G-H)** F-actin-JASP-aged 1 (shades of purple) and **(I-J)** F-actin-JASP-aged 2 (shades of lavender). Both F-actin-PHD and F-actin-PHD-Alexa have only weak density for the inorganic phosphate (P_i_, highlighted by a darker shade of orange), thus, it was not included into the atomic models. Also see Figure 2, Figure 5 and Figure S2, which also include residue labels.

**Movie S3. Binding site of phalloidin and jasplakinolide, Related to Figure 3 and Figure 6**.

Illustration of the phalloidin and JASP (both yellow) binding site consisting of three subunits from both strands. Filament overview and close-up view of the binding site of **(A-B)** F-actin-PHD (shades of blue), **(C-D)** F-actin-PHD-aged (shades of green), **(E-F)** F-actin-PHD-Alexa (shades of turquoise), **(G-H)** F-actin-JASP-aged 1 (shades of purple) and **(I-J)** F-actin-JASP-aged 2 (shades of lavender). The extra density which is present in all phalloidin structures and possibly corresponds to an ion is shown as red mesh in close-up views. Nucleotides are colored in orange. Also see Figure 3, Figure 6, which also include residue labels, and Figure S3.

**Movie S4. Structural changes upon PO4 release in phalloidin-stabilized F-actin. Related to Figure S4**.

Illustration of structural changes between the reconstructions of F-actin-PHD (blue) and F-actin-PHD-aged (green) which can be attributed to the release of P_i_ from the active site. The video is based on a morph of the corresponding electron density maps of the central F-actin subunit. **(A)** Overview of the electron density map of F-actin-PHD. **(B)** Close-up view of the nucleotide binding pocket with focus on Mg^2+^ (green) and P_i_ (orange). In absence of P_i_ (transition from blue to green), Mg^2+^ moves towards the previously occupied position. Moreover, both phosphate binding loop 2 (turquoise, residues 156-159) and switch loop, including the methylated His73, (pink, residues 70-78) move inwards partially closing the empty P_i_ site. **(C)** Tilted view focusing on the inwards movement of phosphate binding loop 2. An additional smaller movement of phosphate binding loop 1 (green, residues 14-16) can be noticed. **(D)** Tilted top view showing the movement of the two phosphate binding loops. The switch loop known to be of importance for phosphate release follows the inwards movement and is furthermore in direct contact with phalloidin (yellow). **(E)** Side view with focus on the switch loop. The proline rich loop (dark red, residues 108-112) also undergoes a minor structural change and is visible in the background. **(F)** Backside view of the complete subunit illustrating the domain movement of SD 1 and SD 2. Upon P_i_ release (transition from blue to green), structural rearrangements within the aforementioned loops are transmitted to connected subunits (primarily SD 2) and eventually cause the downward movement of SD 2 and the adjacent SD 1 which is accompanied by a conformational change of the D-loop and C-terminus. Note that neighboring subunits within one filament might also contribute to the observed structural changes considering the essential intra-strand interactions of the D-loop and C-terminus of two adjacent subunits. Phalloidin (yellow) directly interacts with both the switch (pink) and proline rich loop (dark red). Arg177 was proposed to shuttle P_i_, but does not change its position in this morph. **(G)** Side view showing the downward movement of SD 2 accompanied by refolding of the D-loop and C-terminus. **(H)** Front view illustrating the motion of SD 2 and SD 1, followed by a small movement of SD 3 which is linked to the D-loop of the adjacent subunit through the W-loop (khaki, 165-172), especially Tyr169. For every view the morph of F-actin-PHD (blue, bound ADP and sub stochiometric amounts of P_i_) and F-actin-PHD-aged (green, bound ADP) is looped twice for two different density thresholds each (one optimized for the visualization of P_i_ and the other for Mg^2+^). Throughout **(B-H)** the atomic model F-actin-PHD is displayed for better orientation. An atomic model of P_i_ (not included into the atomic model of F-actin-PHD) is additionally shown in **(B-D)** to guide the identification of the corresponding density. For an overview of the domain organization and localization of loops in the F-actin protomer also see Figure S4.

**Movie S5. Variation along the first principle component of the PC analysis, Related to Figure 7**.

Illustration of the variations within the central F-actin monomer projected onto the first principle component. **(Left panel)** Plot of the first and second principle component (also see Figure 7). The black line highlights the propagation of the model (right panel) along the corresponding principle component. **(Right panel)** Trajectory representing an interpolation between the two most dissimilar structures in the distribution along the first principle component, namely F-actin-ADP and F-actin-JASP-ADP. As the changes are subtle and difficult to appreciate, four different orthogonal views are provided including **A)** front, **B-C)** two side views and **D)** top view. The dominant variation is an upwards rotation of SD 1 and SD 2 accompanied by the reorganization of both the D-loop and C-terminus, similar to what we have reported previously (Merino et al., 2018). Only Cα atoms present in all structures were used for the analysis, thus, part of the D loop and the C terminus were not included and are also missing in the shown model. Both, the D-loop (residues 37-53, blue arrow head) and one connecting loop (126-129, grey arrow head) show high variation. While the change of the D-loop is supported by the electron density maps, changes in the other loop are not meaningful as the corresponding electron density is often fragmented and thus does not allow reliable modelling.

**Movie S6. Variation along the second principle component of the PC analysis, Related to Figure 7**.

Illustration of the variations within the central F-actin monomer projected onto the second principle component. **(Left panel)** Plot of the first and second principle component (also see Figure 7). The black line highlights the propagation of the model (right panel) along the corresponding principle component. **(Right panel)** Trajectory representing an interpolation between the two most dissimilar structures in the distribution along the second principle component, namely F-actin-JASP-aged 1 and F-actin-AMPPNP. As the changes are partially subtle and difficult to appreciate, four different orthogonal views are provided including **A)** front, **B-C)** two side views and **D)** top view. The dominant variation is an rotation or torsion of SD 1 and SD 2 against SD 3 and SD 4. Only Cα atoms present in all structures were used for the analysis, thus, part of the D loop and the C terminus were not included and are also missing in the shown model. The D-loop (residues 37-53, blue arrow head) and two additional loops (126-129, grey arrow head and 23-27, green arrow head) show noticeable variation. The motion of loop 126-129 is however not meaningful as described in the legend of Movie S5.

**Movie S7. Variation along the third principle component of the PC analysis, Related to Figure 7**.

Illustration of the variations within the central F-actin monomer projected onto the third principle component. **(Left panel)** Plot of the first and third principle component (also see Figure 7). The black line highlights the propagation of the model (right panel) along the corresponding principle component. **(Right panel)** Trajectory representing an interpolation between the two most dissimilar structures in the distribution along the third principle component, namely F-actin-PHD-aged and F-actin-ADP-Pi. As the changes are subtle and difficult to appreciate, four different orthogonal views are provided including **A)** front, **B-C)** two side views and **D)** top view. The dominant variation resembles a breathing motion of the complete monomer into all direction. Only Cα atoms present in all structures were used for the analysis, thus, part of the D loop and the C terminus were not included and are also missing in the shown model. The D-loop (residues 37-52) is labeled with a blue arrowhead.

